# Conservation of structure, function and inhibitor binding in UNC-51-like kinase 1 and 2 (ULK1/2)

**DOI:** 10.1101/550111

**Authors:** Apirat Chaikuad, Sebastian E. Koschade, Alexandra Stolz, Katarina Zivkovic, Christian Pohl, Shabnam Shaid, Huiyu Ren, Lester J. Lambert, Nicholas D.P. Cosford, Christian H. Brandts, Stefan Knapp

## Abstract

Autophagy is essential for cellular homeostasis and when deregulated this survival mechanism has been associated with disease development. Inhibition of autophagy initiation by inhibiting the kinase ULK1 has been proposed as a potential cancer therapy. While inhibitors and crystal structures of ULK1 have been reported, little is known about the other closely related kinase ULK2. Here we present the crystal structure of ULK2 in complex with ATP competitive inhibitors. Surprisingly, the ULK2 structure revealed a dimeric assembly reminiscent of dimeric arrangements of auto-activating kinases suggesting a role for this association in ULK activation. Screening of a kinase focused library of pre-clinical and clinical compounds revealed several potent ULK1/2 inhibitors and good correlation of inhibitor binding behavior with both ULK kinases. Aurora A was identified as a major off-target of currently used ULK1 inhibitors. Autophagic flux assays demonstrated that this off-target activity by strongly inducing autophagy in different cellular systems conferred an additional layer of complexity in the interpretation of cellular data. The data presented here provides structural models and chemical starting points for the development of ULK1/2 dual inhibitors with improved selectivity for future exploitation of autophagy inhibition.

## INTRODUCTION

Autophagy is an essential mechanism for the survival of eukaryotic cells upon low nutrient availability and prolonged starvation, and at the basal level serves a recycling role for cellular components through degradation of a wide range of substrates from proteins to organelles required for maintaining cellular homeostasis^1–2^. This catabolic process constitutes a multi-step activation and maturation mechanism involving the assembly of large signalling complexes of more than 40 conserved proteins^3–5^. During autophagy, cytoplasmic contents are sequestered within the developing double-membraned phagophore, from which the completed autophagosome then fuses with the lysosome leading to degradation of the contents with subsequent release of free amino acids and other byproducts. In the past few decades, the importance of this process in human health has been demonstrated by a number of studies, revealing two contradictory faces of autophagy. While exerting a protective role against cellular stress^6^ and development of a number of diseases including cancer, neurodegenerative and hyperinflammatory disorder^7–11^, autophagy appears to strongly contribute to the progression of cancer, such as KRAS-driven tumors, to survive a nutrient-deprived microenvironment and therapeutic stress^12–19^. Because of the latter, autophagy inhibition may provide a strategy to inhibit tumor cell growth and to improve efficacy of cancer treatments^20–21^. However, the potential benefits of the therapeutic targeting of autophagy remains controversial due to conflicting context-dependent roles of this process in cancer development and progression^22–24^, and therefore there is a need for a better understanding on the complexity of its regulation.

Initiation of autophagy requires the formation of an early regulatory complex centered on Unc-51 like autophagy activating kinase 1 (ULK1), a serine/threonine kinase homolog of yeast ATG1 (autophagy-related protein 1), that recruits its binding partners ATG13 (autophagy-related protein 13), RB1CC1 (RB1-inducible coiled-coil protein 1 known also as FIP200) and ATG101 (autophagy-related protein 101)^25–27^. mTOR inhibition results in activation of early autophagy, a process that is counteracted by the metabolic sensor kinase AMPK linking these two key pathways regulating cellular metabolism. In humans, ULK1 cooperates in regulating autophagosome assembly with a closely-related homolog, ULK2, both of which share a high degree of conservation of the domain architecture including an N-terminal catalytic kinase, extensive middle linker and C-terminal domain^28^. These two kinases have an overlapping function in autophagy induction under nutrient deprivation^29–30^. As a consequence, complete knock-outs of both kinases in mouse embryonic fibroblasts is required for inhibition of autophagy^30–32^. Nevertheless, some non-overlapping roles of ULK2 has been demonstrated recently, which might potentially relate to context dependent requirements and the genetic background of specific cell types^33–35^.

With essential roles in diseases^36–40^, ULK1 has been identified as an attractive target for inhibition of the autophagy pathway. Crystal structures of ULK1 provide insights into the druggable pockets of this kinase^41^, leading to the development of several inhibitors, including SBI-0206965^42^, MRT67307, MRT68921^43^ and recently ULK-101^44^. The efficacies of these small molecules towards autophagy suppression has been demonstrated, providing an initial proof of concept for this kinase as a potential chemotherapeutic target. However, while published inhibitors are potent inhibitors of ULK1, currently available inhibitors are not entirely selective for ULK1. Furthermore, to date, little is known regarding the role of the closely related isoform ULK2, which according to its sequence and domain conservation would also need to be targeted for the pharmacological modulation of autophagy initiation. The lack of information regarding the structural, biochemical and biological properties of ULK2 therefore may limit our understanding on potential beneficial effects and applications of the use of these ULK1 inhibitors for suppression of autophagy.

In this study, we report the first structure of ULK2, revealing not only similarity of its kinase domain and ligand binding properties to that of the family-related ULK1, but also an interesting activation segment domain exchange dimerization that potentially represents a structural mechanism for autophosphorylation and activation of these two kinases. Screening against a diverse set of kinase inhibitors showed similar inhibitor binding properties of both ULK isoforms and demonstrated unexpectedly potent hits of diverse inhibitors developed for other kinases. The binding of these compounds is consistent with the flexible nature of the ULK1/2 kinase pockets whereas notably Aurora kinase activity seems a liability of current ULK1 tool compounds and many of the identified inhibitors in our screen. Structural and functional consequences of this unexpected SAR (structure activity relationship) similarity are discussed.

## EXPERIMENTAL SECTION

### Protein expression and purification

All ULK1 (1-283) and ULK2 (1-276) kinase domains were recombinantly expressed as fusion proteins incorporating either His_6_ or His_6_-Sumo tags at the N-termini. Briefly, *E. coli* cultured in TB media was initially grown at 37 °C until reaching the OD_600_ of 1.6-1.8, was then subsequently cooled to 18 °C and at OD_600_ of 2.6-3.0 was induced with 0.5 mM IPTG overnight. Harvested cells were lysed by sonication, and the proteins were purified using Ni^2^+-affinity chromatography. The eluted proteins were buffer exchanged into 30 mM Tris, pH 7.5, 300 mM NaCl, 0.5 mM TCEP and 10% glycerol, and their expression tags was cleaved using SENP1 or TEV. If necessary autophosphorylation was performed during proteolysis by supplementing ATP and MgCl_2_ at 10- and 20-fold molar excess, respectively. The cleaved proteins were passed through Ni^2+^ beads, and further purified by size exclusion chromatography. The pure proteins in 25 mM Tris, pH 7.5, 100 mM NaCl, 10% glycerol were stored in −80 °C.

### Crystallization and structure determination

The recombinant ULK1 and ULK2 at ~10-12 mg/ml were mixed with inhibitors at ~2-3-fold molar excess prior to crystallization using the sitting drop vapor diffusion method at 20 °C. Crystallization conditions are summarized in Supplementary table 2. Viable crystals were cryo-protected using mother liquor supplemented with 25% glycerol or 22% ethylene glycol, flash frozen in liquid nitrogen, and tested for their X-ray diffraction quality at Bessy II. Diffraction data were collected at Swiss Light Source or Diamond and was processed and scaled with iMosflm^45^ or XDS^46^ and aimless^47^, respectively. Initial structure solutions were obtained by molecular replacement using Phaser^48^ and the coordinates of previously published ULK1^41^. Iterative cycles of manual model rebuilding alternated with structure refinement were performed in COOT^49^ and REFMAC^50^, respectively. Final structures were verified for their geometric correctness using molprobity^51^. The data collection and refinement statistics are summarized in Supplementary table 2.

### Thermal shift assays

Recombinant kinase domains of ULK1, ULK2 and Aurora A at 2 μM in 10 mM HEPES, pH 7.5 and 500 mM NaCl were mixed with 10 μM inhibitors. Temperature-dependent protein unfolding profiles were measured using a Real-Time PCR Mx3005p machine (Stratagene). The data evaluation and melting temperature calculation were performed as described previously^52^’^53^.

### Isothermal calorimetry

All isothermal calorimetry (ITC) experiments were performed on NanoITC (TA instrument) at 25 °C in the buffer containing 25 mM Tris, pH 7.5, 200 mM NaCl, 0. 5 mM TCEP and 10% glycerol. The protein at ~0.12 mM was titrated into the reaction cell containing the inhibitors. The heat of binding was integrated, corrected and fitted to an independent single binding site model based on the manufacture protocol, from which thermodynamics parameters (ΔH and TΔS), equilibrium association and dissociation constants (Ka and K_D_) and stoichiometry (n) were calculated.

### Autophagic flux assays in RPE1

RPE1 cells (1500 cells/well in 50 μl DMEM/F12, 10% FBS, 1% penicillin/streptamycin) stably expressing the GFP-LC3-RFP-LC3ΔC autophagic flux reporter^54^ were seeded in black, optically clear bottom 384-well plates (Nunc) and grown for ~18 hours. Plates were subsequently placed and monitored in the IncuCyte^®^ (Sartorius). After the first scan (0 hr), additional 50 μl media containing compounds at 2x final concentration were added. Cells were scanned continuously during the measuring time window for phase contrast and fluoresces to obtain information about confluency and autophagy flux, respectively.

### Autophagic flux assays in leukemia cell lines

Human AML cell lines MV4-11 and MOLM-14 (purchased from Leibniz-Institut DSMZ-Deutsche Sammlung von Mikroorganismen und Zellkulturen GmbH [DSZMZ]) stably expressing a GFP-LC3B-mCherry flux reporter^54^ after lentiviral transduction similarly as previously described^61^ were cultured in RPMI 1640 Medium (Gibco) supplemented with 10% fetal bovine serum (FBS) (Sigma-Aldrich) and 1% penicillin/streptamycin. Cell lines were cultured at 37 °C with 5% CO2 in a humidified Heracell 150i incubator (Thermo Fisher Scientific) and regularly confirmed by PCR to be free of mycoplasms. For dose-response measurements, cells were washed once with PBS (Gibco) on the day prior to the experiment and cultured in fresh medium at a density of 0.5×10^6^ cells/ml. On the next day, cells were washed again, resuspended in fresh medium, seeded at a density of 0.5×10^6^ cells/ml and treated with individual compounds or DMSO as a vehicle control. Fluorescence mCherry (G610/20) and eGFP (B530/30) signal intensities were measured by flow cytometry on a BD LSRFortessa (BD Biosciences) flow cytometer after 8h and 24h following treatment. At least 10,000 events were measured for each sample. Autophagic flux was quantified by ratiometric single-cell eGFP/mCherry values, aggregated per sample by the arithmetic mean, and normalized to DMSO vehicle controls. Experiments were performed in duplicates.

### ADP-Glo assay

For the ULK1 kinase assay, the 5 uL kinase reaction was performed with 2 ug/mL recombinant human ULK1 protein (1-649, SignalChem #U01-11G) and 80 ug/mL myelin basic protein (MBP, Sigma-Aldrich #M1891) in the presence of 25 uM ATP (Sigma-Aldrich A7699). For ULK2 kinase assay, the 5 uL reaction was performed with 4 ug/mL recombinant human ULK2 protein (1-478, SignalChem #U02-11G) and 80 ug/mL MBP in the presence of 25 uM ATP. Compounds were tested in triplicate in a 16-dose IC_50_ mode with 3-fold serial dilution and a starting dose of 30 uM. Staurosporine, a non-selective protein kinase inhibitor, was used in the assay as a positive control. Three separate experiments were carried out.

## RESULTS AND DISCUSSION

### ULK1 and ULK2 share a conserved overall kinase structure

Sequence analysis of ULK1 and ULK2 revealed that the kinase domain of both proteins shared a high sequence identity of ~75%, and apart from an N-terminal extra 7-amino acid extension in ULK1 all catalytic elements essential for kinase activities were well conserved (Supplementary Figure 1). To provide structural models, we crystallized both kinases and determined the high-resolution crystal structures of ULK1 harboring the E37A, K38A surface entropic mutation doubly phosphorylated at S87 and the activation loop T180 (Figure 1A) and the activation loop T173D ULK2 mutant (Figure 1B). Both ULK1 and ULK2 were highly resembled in overall topology, adopting the typical bilobal kinase architecture similar to the previously published ULK1 structural model^41^ (Figure 1 and Supplementary Figure 1). However, two distinct structural differences were observed at the regulatory helix αC and the activation segment. In ULK1 harboring a phosphorylated activation loop at T180 these two regulatory elements assumed an active conformation, similar to the ULK1 structure reported previously^41^, while the ULK2 structure displayed a distorted αC and an elongated activation segment, likely depicting an inactive conformation despite containing the activation loop T173D phosphomimetic mutation. Other less dramatic conformational differences were apparent for the flexible β6-β7 insertion loop (interlobe loop) that shared low sequence identity and the β4-β5 loop of which the unexpected non-regulatory autophosphorylation at S87 in ULK1 caused small perturbation on the N-lobe β-sheet integrity. Nonetheless, apart from these alterations both structures superimposed well, demonstrating a high structural conservation between these two key autophagy kinases (rmsd of ~1.3 Å for 220 Cα atoms, excluding the αC helix and activation segment).

**Figure 1.**
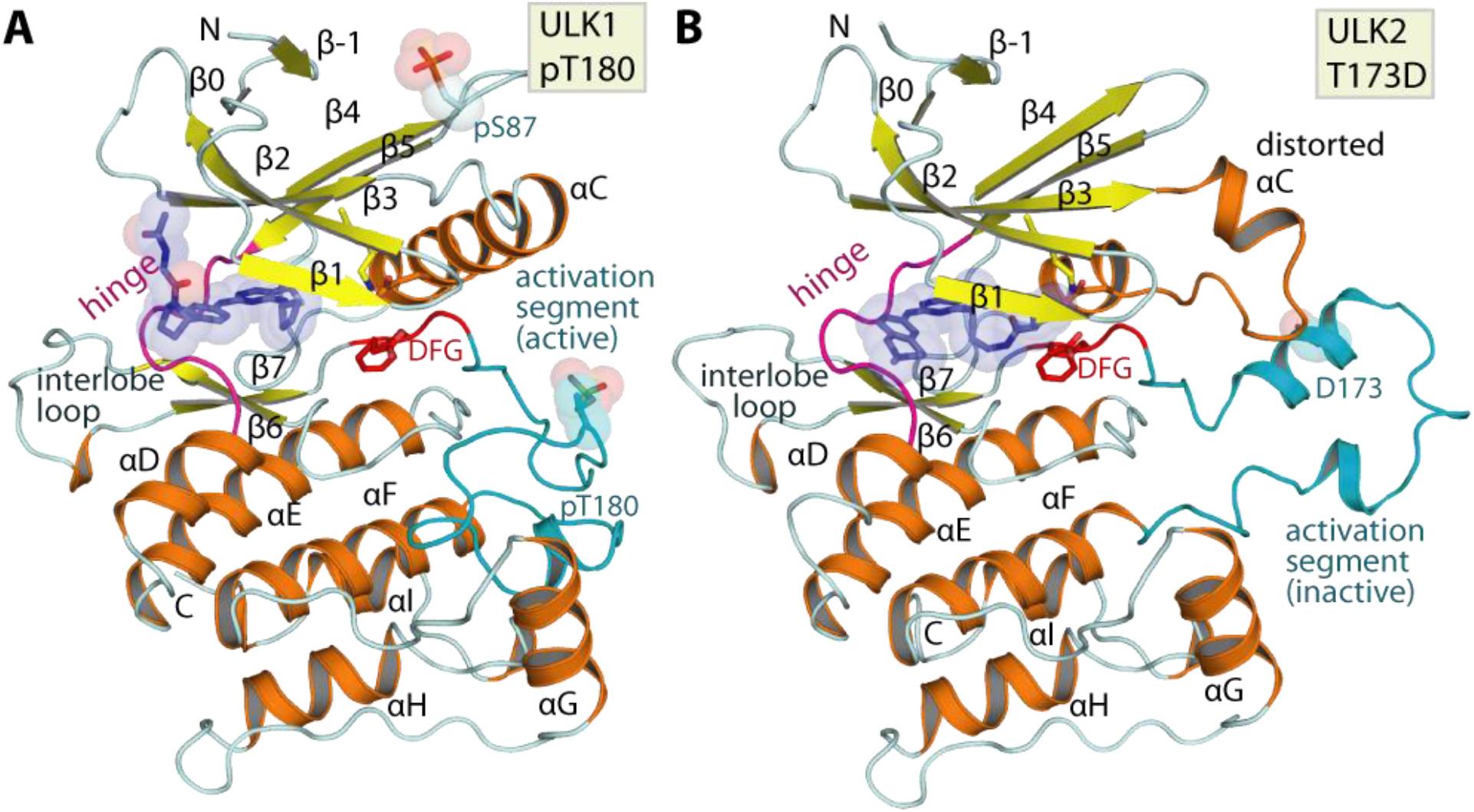
Overall structure of ULK1 and ULK2. A) Crystal structure of PF-03814735-complexed ULK1, doubly phosphorylated at S87 and T180, depicting an active conformation of the kinase. B) Structure of phosphomimetic T173D ULK2 mutant in complexed with MRT68921 surprisingly revealed an inactive kinase conformation depicted by distorted αC and the inactive conformation of the activation segment. Main structural elements are labelled and co-crystallized inhibitors are shown in blue stick representation with semitransparent surfaces.

### Activation loop exchange enables dimeric assembly in ULK2 structure

Analyses of the crystal structures revealed another unexpected feature in ULK2. The ULK2 catalytic domain engaged in a dimeric assembly, which was in contrast to the typically monomeric ULK1. The two protomers of the ULK2 kinase domain associated in a face-to-face fashion with the αC and the activation segment packed at the two-fold dimeric interface, explaining the unusual conformations of these two structural elements (Figure 2A and Supplementary Figure 2). A striking feature in the dimer was the exchange of the activation segment, a configuration involving a protrusion of the segment from one subunit into the substrate binding groove fenced by the αF and αG of the neighboring subunit in the pair. Despite a few direct inter-molecular interactions formed (Figure 2B), this extended conformation covers a large interface contact area indicating a potential driver for this oligomerization.

**Figure 2.**
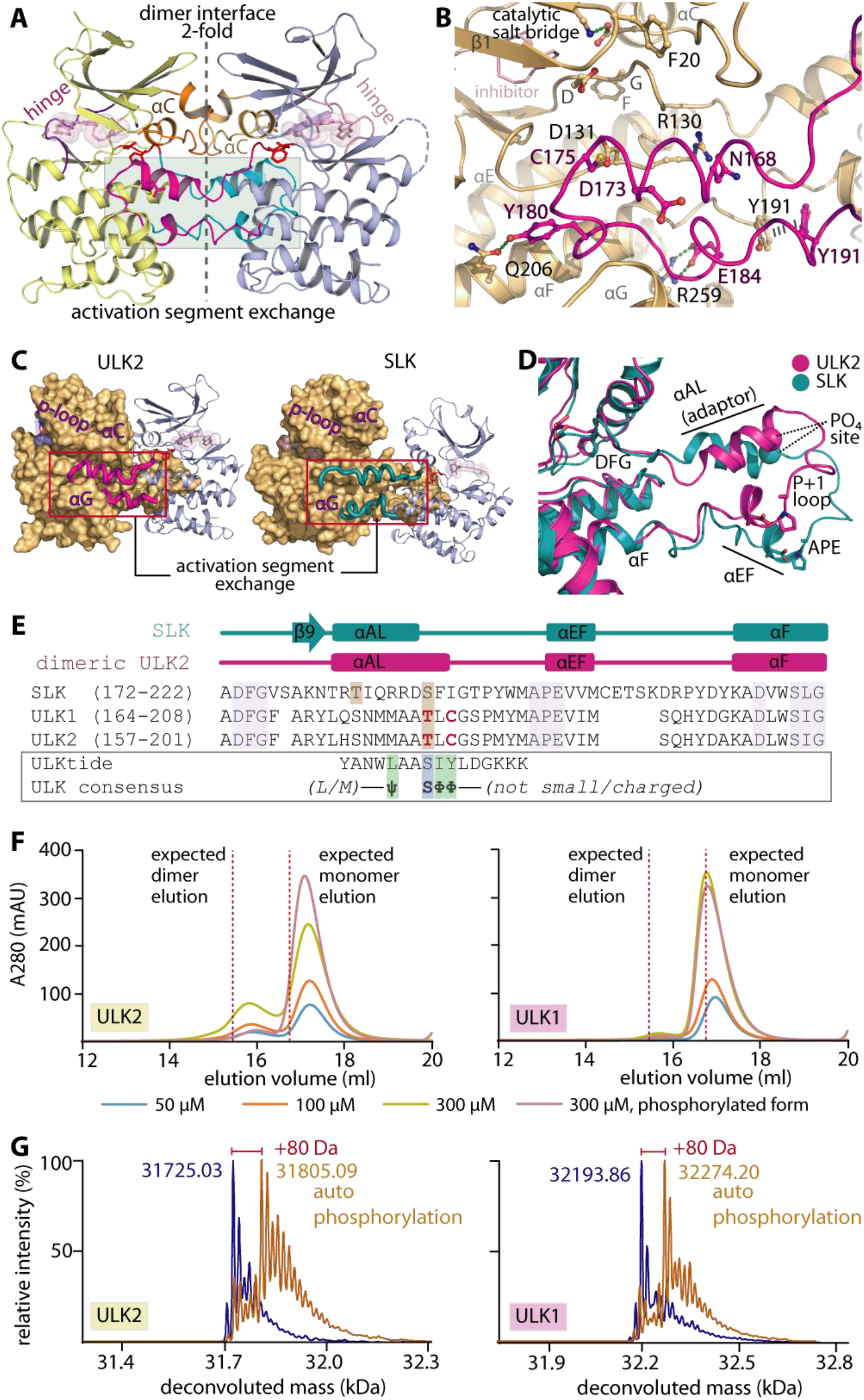
Domain-exchanged activation segment dimerization in ULK1/2. A) Face-to-face dimeric assembly in ULK2-MRT68921 structure showing the packing of αC and domain-exchanged activation segment at the two-fold interface. B) Detailed interactions between the protruding activation segment of one subunit and the active site of the neighboring kinase in the dimer. C) ULK2 and SLK shares similar domain-exchanged activation segment configuration for their dimerization. D) Superimposition of activation segments of ULK2 and SLK reveals a highly similar conformation. E) Structure-based sequence alignment shows that the threonine phosphorylation sites of ULK1 and ULK2 are well aligned with the S189 residue of SLK that is known to be an autophosphorylation site (brown boxes). These threonine residues are located within the region of the activation segments that lacks consensus substrate sequence, for which the residues that violate the pattern are highlighted in bold, red letters. F) Gel filtration elution profiles of wild-type ULK1 and ULK2 demonstrate potentially transient existence of dimers in solution for non-phosphorylated proteins, which decreases in the phosphorylated kinases. G) Mass spectrometry analyses reveals an increase in ~80-dalton mass upon incubating ULK1/2 with ATP and MgCl2, indicating the kinase autophosphorylation.

We were interested whether the dimer observed in the ULK2 crystal structure is physiologically relevant or may serve a regulatory function. Comparative structural analyses revealed that this face-to-face, domain-exchanged activation segment dimerization in ULK2 highly resembled conformations observed previously in other kinases, such as SLK, LOK, DAPK3 and CHK2^55–56^ (Figure 2C-E and Supplementary figure 2). Since the dimeric exchanged activation segment arrangement has been proposed as a mechanism for *trans* activation by autophosphorylation of the activation segment at non-consensus sites^55, 57^, we therefore postulated a similar role for this conformation in ULK1/2. This hypothesis was supported by i) a non-consensus sequence present in the ULK1/2 activation segment, exemplified by T180/173 and C182/175 that did not match the preferred serine and bulky hydrophobic amino acids flanking the phosphorylation sites in consensus substrate sequences^42, 58^ (Figure 2E), and ii) the presence of the activation loop helix (αAL) adaptor which has been shown to be necessary for positioning the activation segment threonine into the kinase substrate site (T180 and T173 in ULK1 and ULK2, respectively, equivalent to S189 in SLK^55^) (Figure 2B-E). In addition, although this self-association stabilized in the crystals is typically a transient property in solution^55^, we were able to detect dimerization of ULK2 in analytical gel filtration for both wild-type, non-phosphorylated ULK2 and, to a lesser extent, wild-type ULK1 (Figure 2F). The functional consequence of this interaction was demonstrated by the ability of the two kinases to autophosphorylate upon incubating with ATP-MgCl_2_ (Figure 2G), from which the location of the phosphothreonine at the corresponding sites in ULK1/2 was confirmed by tryptic digestion mass spectrometry (data not shown). This cumulative evidence potentially suggested therefore that ULK1/2 may possess an ability for (transient) dimeric assembly, and utilize the shared activation segment domain-exchanged mechanism for *trans* autophosphorylation of their specific activation loop threonine residues for kinase activation. Nevertheless, the roles of the distorted, inactive αC-helix conformation observed in the ULK2 structures (Figure 1B) in its trans-activation mechanism remain unclear.

### Conservation of inhibitor binding in ULK1 and ULK2

The high sequence and structural similarity between ULK1 and ULK2 prompted us to next investigate whether these kinases also share similar inhibitor binding properties. A set of 384 known kinase inhibitors, which included also some previously developed ULK1 inhibitors such as SBI-0206965^42^, MRT67307 and MRT68921^43^, were screened against the kinases using thermal stability-shift assays^52^. As expected, a good correlation of the temperature shift (ΔTm) profiles was observed across this set of compounds for the ULK1 and ULK2 catalytic domains indicating that both kinases exhibit similar inhibitor binding behavior (Figure 3A). Among the three ULK1 inhibitors, MRT68921 results in the highest Tm shift for both kinases (ΔTm of 12.1 and 14.8 °C for ULK1 and ULK2, respectively), while the others show only moderate stabilization with similar ΔTm values in a range of 6.5-9.8 °C (Figure 3A and Supplementary Table 1).

**Figure 3.**
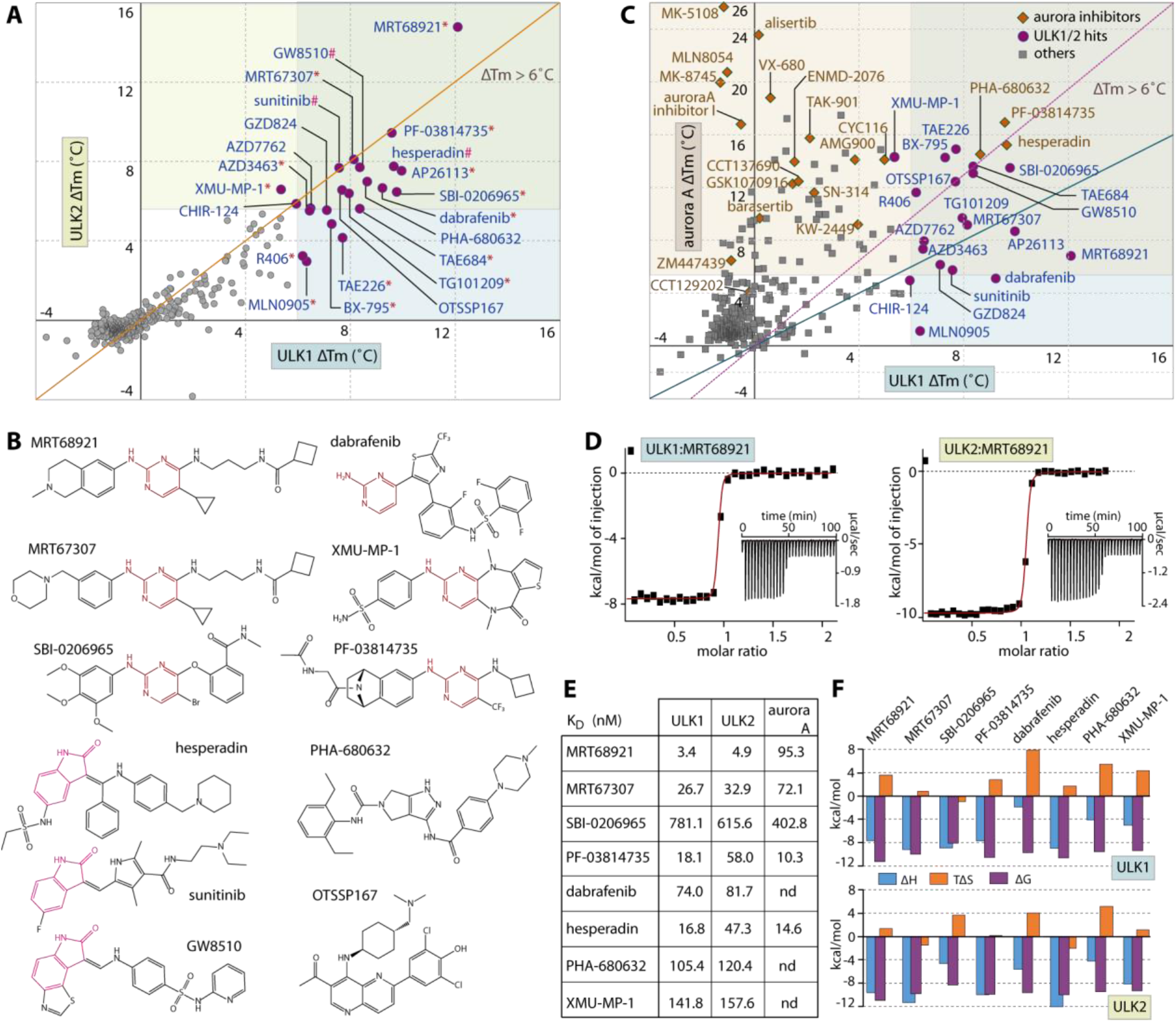
Inhibitor screening for ULK1 and ULK2. A) The plot of Tm shifts (ΔTm) of ULK1 against ULK2 for the set of 384 known kinase inhibitors demonstrates good correlation in inhibitor binding behavior for both kinases (orange, diagonal line indicates equal points on both axes, and the data are also summarized in Supplementary Table 1). The 22 hits with ΔTm more than 6 °C were grouped into three structural classes: 2-aminopyrimidine (indicated by *), 2-oxindole (#) and others (no symbol), of which example chemical structures of some inhibitors are shown in (B). C) Analyses of the ΔTm of ULK1 and Aurora A reveals common cross-activities of nearly all of ULK1 hits on Aurora A kinase, but not vice versa. The green diagonal line indicates equivalence of both values, while the magenta, dotted line intersects the highest and lowest points of both axes on the shown scales. D) Example ITC binding data for the interactions between MRT68921 with ULK1 and ULK2. The normalized heat of binding with the single-site binding fits (red line) are shown, while the raw isotherms of titration heat are displayed as insets. The binding affinities and the thermodynamics signatures from ITC for the selected set of compounds are shown in E) and F), respectively.

Interestingly, there were 18 additional, structurally diverse compounds that also exhibited strong Tm shifts with similar levels to that of the ULK1 inhibitor SBI-0206965 and MRT67307 as indicated by ΔTm of > 6 °C, typically suggesting a binding affinity in sub-micromolar range^53^ (Figure 3A and Supplementary Table 1). Structural analyses reveals that most of these compounds share two generic hinge binding motifs of either 2-aminopyrimidine or 2-oxindole, yet other scaffolds are also evident (Figure 3B). Nonetheless, the large number of hits from this set was rather unexpected considering the structure-activity relationships that presumably optimized them for their specific kinase targets. For example, the activities of a number of specific Aurora kinase inhibitors, such as hesperidin, PF-03814735 and PHA-680632, towards ULK1/2 were surprising considering that they belong to two distinct kinase subfamilies. To investigate whether ULK1/2 and Aurora A might exhibit similar preference towards certain inhibitors, we compared the Tm shifts of ULK1 and Aurora A, and observed good correlations for some compound classes (Figure 3C). Nearly all inhibitors that bound ULK1 also interacted strongly with Aurora A, and this applied also to the ULK tool inhibitors SBI-0206965, MRT67307 and MRT68921. However, this trend was not observed for all Aurora A inhibitors that have been screened, suggesting inhibitor class dependent cross-reactivity. For example, the Aurora A inhibitor alisertib, a pyrimido benzazepine, that enters clinical trials was inactive against ULK1 and ULK2.

To validate the results from the Tm shift assays, we performed isothermal calorimetry (ITC) to measure binding affinities of some ULK1/2 hits in solution. All selected inhibitors showed K_D_ values for ULK1, ULK2 and Aurora A in the low nanomolar range in agreement with the observed Tm shifts and the measured *in vitro* IC_50_ values (Figure 3D-F and Table 1).

**Table 1.** IC_50_ values for current ULK1 and ULK2 inhibitors measured by ADP-Glo assays. The assay data represent averaged values from three independent experiments.

|  | ULK1 |  | ULK2 |  |
| --- | --- | --- | --- | --- |
| | Average $IC_{50}$ (M) | SEM (M) | Average $IC_{50}$ (M) | SEM (M) |
| MRT67307 | 3.82E-08 | 7.39E-09 | 9.23E-08 | 2.66E-08 |
| MRT68921 | 1.70E-08 | 4.52E-09 | 2.08E-08 | 4.58E-09 |
| SBI-0206965 | 3.06E-07 | 1.59E-07 | 3.88E-06 | 9.02E-07 |

Increasing affinities corresponded well with higher ΔTm values. MRT68921, for example, was the most potent inhibitor in the Tm assay with a K_D_ of 3.4 and 4.9 nM for ULK1 and ULK2, respectively and the affinities of ~17-160 nM for the others correlated well with their lower ΔTm values in the 6-10 °C region (Figure 3E). An exception was noted for the moderate affinities for SBI-0206965, which correlated well with the ΔTm values and matched the reported IC_50_ value for ULK2^42^. However, our *in vitro* K_D_ for ULK1 was ~7-fold smaller compared to the previously reported potency on ULK1 (IC_50_ of 108 nM^42^), yet consistent with the micromolar activities recently reported in cellular systems^44^. Nonetheless, regardless of their potencies most of these typical type-I inhibitors exhibited similarly entropically favorable thermodynamic signature for their binding in both kinases (Figure 3F).

### Structural insights into the common binding pockets of ULK1 and ULK2

To provide structural information of the inhibitor binding modes, we determined high resolution crystal structures of four kinase-inhibitor complexes, including ULK1 with PF-03814735, and ULK2 with MRT68921, MRT67307 and hesperadin (Figure 4A). All inhibitors exhibited typical type-I interactions with the canonical hydrogen bonds formed by the 2-aminopyrimidine and 2-oxindole scaffolds with the hinge backbones. However, small differences were noted in their binding positions such as the deep protrusion of 2-oxindole towards the back pocket targeting the residue at the gatekeeper +1 (GK+1) position, which was in contrast to interactions formed by 2-aminopyrimidine that was shifted outward to interact instead with the gatekeeper +3 position (GK+3) residue (Figure 4B). In comparison, all of these positions have been exploited previously for an accommodation of the alternative 3-aminopyrazole hinge-binding motif^41^. These data suggest therefore a potential capacity for ULK1/2 in accepting diverse type-I hinge binding scaffolds which is supported by the high hit rates in the small focused library screen.

**Figure 4.**
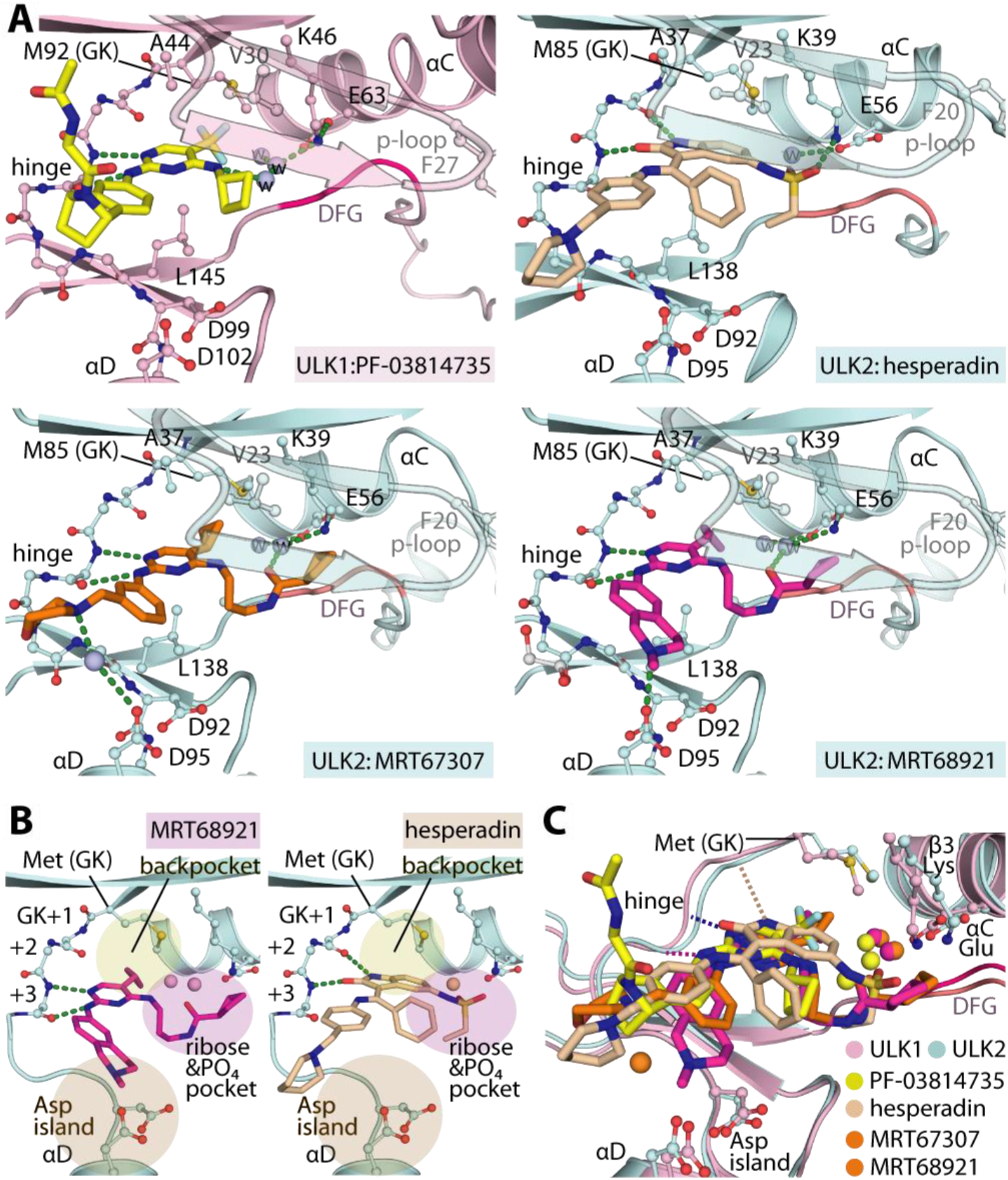
Crystal structures of ULK1/2-inhibitor complexes. A) Detailed interactions between ULK1 and PF-03814735 and ULK2 with hesperadin, MRT67307 and MRT68921. Bound water molecules are shown in spheres. B) Schematic illustrations of the binding pockets of the 2-aminopyrimidine-based MRT68921 and 2-oxindole-based hesperadin revealing different interactions of the hinge binding motifs. Three cavities for accommodation of the inhibitors in ULK1/2 are highlighted. C) Superimposition of ULK1/2-inhibitor complexes demonstrated highly similar binding sites in both ULK kinases and also the commonly occupied space by diverse inhibitors.

Detailed analyses of the inhibitor-complexed structures provided further insights into the characteristics of the ULK1/2 binding sites (Figure 4A-C). First, we observed some degree of flexibility of the methionine gatekeeper (M92 and M85 in ULK1 and 2, respectively), suggesting a certain degree of plasticity of the back pocket when accommodating bulky hydrophobic moieties, such as cyclopropane in MRT67307 and MRT68921, trifluoromethyl of PF-03814735, or even bulkier oxindole of hesperidin as well as iodide and cyclobutane reported to occupy this position previously^41, 59^. Secondly, the binding of all inhibitors induced an ‘outward’ conformation of the phenylalanine at the tip of the P-loop (F27 and F20 in ULK1 and 2, respectively), creating an unusually large binding pocket within the ribose and phosphate binding pocket. Evidently, this cavity was exploited by diverse decorating groups of the co-crystallized inhibitors which filled different space leading to different bound-water patterns and engaging slightly different interactions with the kinases. (Figure 4A and 4C). A large pocket was noted in the solvent-exposed region adjacent to the hinge, however the interactions of most inhibitors in this region were rather limited. Interestingly, we observed a unique trajectory of the benzopiperidine of MRT68921 towards the negatively-charged, aspartate rich region in proximity to the αD, leading to a direct contact with D95 (Figure 4B). This distinct interaction might explain the ~8-fold increase in affinity of MRT68921 in comparison to the highly-related MRT67307.

Overall, these structures present unusually large binding pockets in ULK1 and ULK2 suggesting a high degree of plasticity of these kinase catalytic domains, which was in agreement with the unexpectedly large number of diverse, highly-decorated hits identified from the screened kinase inhibitor set. In addition, both kinases employ a similar set of residues that line the binding sites, supporting the close correspondence of their inhibitor binding behaviors (Figure 4A and 4C). Such undistinguishable characteristics of the pockets shared between ULK1 and ULK2 suggest therefore that both kinases can be efficiently co-targeted by an inhibitor with highly similar interactions and binding properties which is supported by similar affinities of known ULK1 inhibitors and the large number of identical hits identified. As efficient inhibition of both ULK1 and ULK2 is required for efficient inhibition of autophagy, our structural data suggests that selective dual-targeting chemical probes can be developed for these two kinases.

### Modulation of Autophagy by current ULK inhibitors

Having demonstrated high similarity of inhibitor binding characteristics in ULK1 and ULK2, the current ULK1 tool compounds may possess sufficient favorable property for achieving inhibition of autophagy at initiation through these kinases. However, cross-activities on off-targets, as demonstrated for example on Aurora A (Figure 3), could lead to more complex effects on autophagy modulation. To assess this, we exploited an LC3-based GFP/mCherry or GFP/RFP fluorescence assay to measure autophagic flux^54^ following treatment with three ULK1 inhibitors, SBI-0206965, MRT67307 and MRT68921, and the highly selective Aurora inhibitor alisertib (Figure 5A). First, using a RPE cell model^60^ under conditions of normal growth media, we observed surprisingly that all inhibitors moderately induced autophagy as observed by the decrease of GFP/RFP ratio throughout the measured time course (Figure 5B-C). Interestingly, all four inhibitors shared a concentration-dependent autophagy induction pattern to that of alisertib (Figure 5C). However, differences in autophagic flux were noticed for the most potent ULK1/2 inhibitor MRT68921, of which the induction effects were less pronounced, and reversed at higher inhibitor concentration which might potentially be due to the anticipated ULK1/2 inhibition.

**Figure 5.**
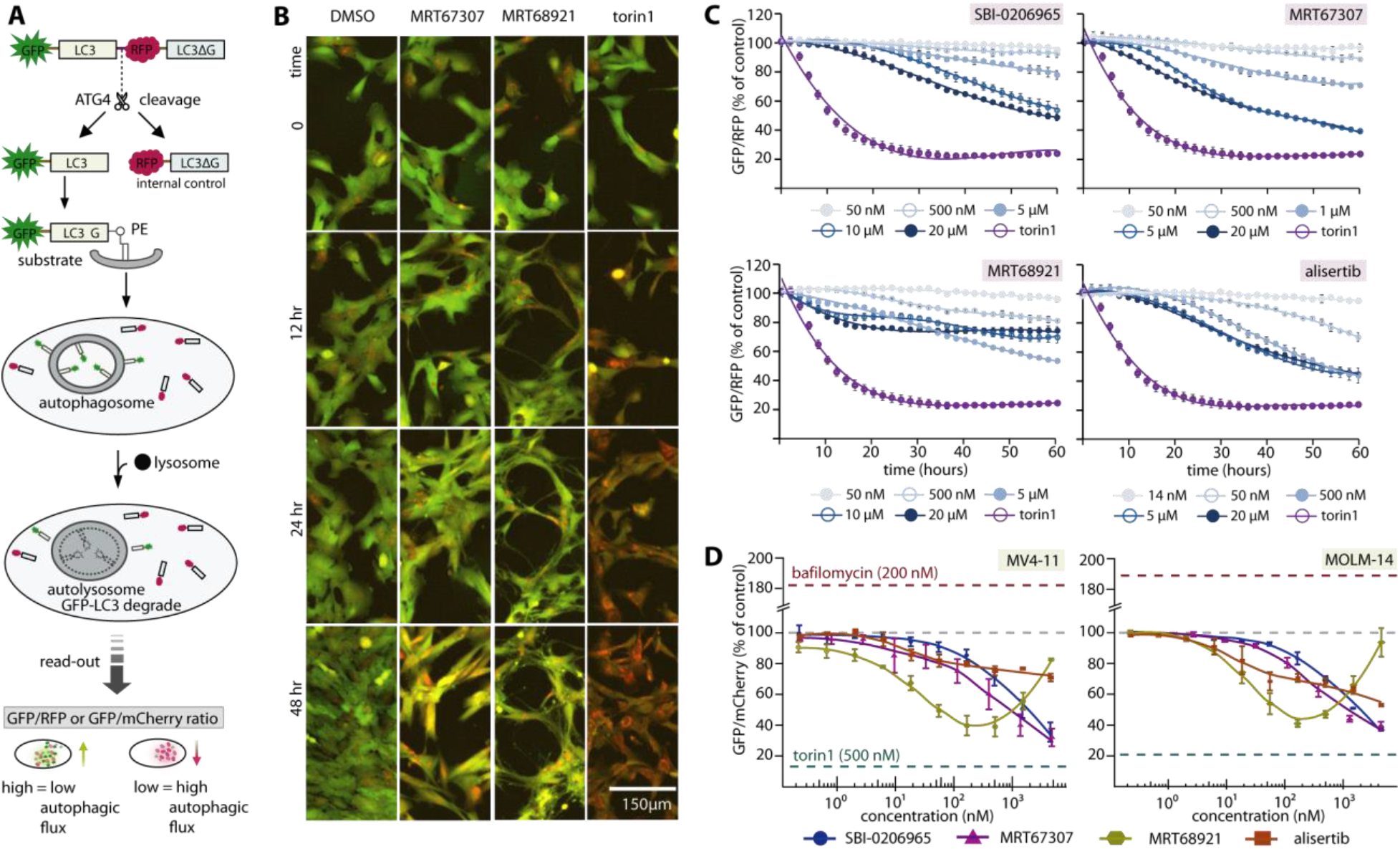
Complex cellular effects of ULK1/2 inhibitors on autophagy modulation due to off-target effects. A) Schematic diagram illustrates LC3-based GFP/RFP fluorescence autophagy flux assays used in RPE1 experiment. A similar assay scheme was used in leukemia cell lines, albeit with an omission of C-terminal LC3ΔG from the GFP-LC3B-mCherry probe. The read-out GFP/RFP or GFP/mCherry ratio from high to low confers low to high autophagy flux levels, respectively. B) Examples of RPE1 cells at different time point in the autophagic flux assays. Fluorescent images (overlay green and red) of RPE1 cells stably expressing the autophagy flux reporter GFP-LC3-RFP-LC3ΔC. Cells were treated for the indicated time with 1% DMSO, ULK1 inhibitor MRT67307 and MRT68921 at 5 μM, and 250 nM torin1, respectively. A drop in GFP (green channel) indicates increased autophagy flux. C) Time-dependent progression of autophagic flux in RPE1 cells of three ULK1 inhibitors and selective Aurora A inhibitor alisertib at various concentrations (torin1 used as control at 250 nM). D) Autophagic flux measured at 24-hour time point reveals dose-response of autophagic flux of the same set of inhibitors in leukemia cell line MV4-11 and MOLM-14. For both (C) and (D), mean of repeated experiment with standard error of the mean (SEM) are shown.

We next performed these assays in two leukemia cell lines, MV4-11 and MOLM-14 (Figure 5D and Supplementary figure 3). Consistently, we observed autophagy induction for SBI-0206965, MRT67307 and alisertib in a similar dose-dependent manner as seen in RPE1 cells. Interestingly, MRT68921 exhibited once again a unique pattern among the tested inhibitors. While causing initial induction of autophagy at lower concentrations, a reverse of fluorescence ratios was evident at higher concentration of >100 nM, suggesting autophagy inhibition at higher compound concentration.

Overall, the concentration- and time-dependent autophagic flux profiles of the tested inhibitors likely indicates that despite potently inhibiting both ULK1 and ULK2 their off-target activities, such as Aurora A inhibition, induced autophagy, most likely caused by cytotoxicity or cellular stress. This clearly complicates readouts on autophagic flux. The macroautophagy induction constitutes a cytoprotective survival process observed in several cancer entities upon cytotoxic treatment^13, 15^. For mechanistic studies, development of more selective ULK1/2 inhibitors with minimal off-target effects is needed for determining the value of ULK1/2 inhibition as a therapeutic target in a definitive manner. Nevertheless, from a therapeutic standpoint, these ULK1 inhibitors, which lead to both cytotoxicity (through Aurora A inhibition, among others) and to inhibition of autophagy as a survival pathway (through ULK1/2 inhibition), may be a novel and effective approach in cancer therapy.

## Conclusion

Autophagy is a complex cellular process that is regulated on many levels and which has been implicated in the development of a large diversity of diseases. In order to explore the potential of this process for the development of new treatment strategies, we presented here several high resolution structural models of the key autophagy inducing kinases ULK1 and ULK2. These structures can now be used for the rational design of highly potent and selective chemical ULK1/2 probes for functional cellular studies. Screening a diverse kinase focused library we identified diverse chemical starting points for inhibitor development. In addition, we identified Aurora A kinase as a major off-target of currently used and available ULK1/2 inhibitors. Autophagic flux assays revealed that Aurora A inhibition by selective Aurora A inhibitors strongly induced autophagy counteracting the inhibitory effects expected by ULK1/2 inhibition. At higher compound concentration however, inhibitory effects on autophagic flux was observed for the most potent ULK1/2 inhibitors. The data demonstrate that for more detailed mechanistic studies, ULK1/2 inhibitors with improved selectivity would be desirable, which forms a current research focus in our laboratories.

## Supporting Information

The supporting information includes figures and tables.

## Author Contributions

AC, CHB and SK designed research. AC performed crystallographic, biochemical and inhibitor binding study. Cellular inhibition study was performed by SEM and SS for AML cell lines, and AS, KZ and CP for RPE1 cells. HR, LJL and NDPC performed ADP-Glo assays. AC and SK wrote the manuscript, which was approved by all authors.

## Notes

The coordinates and structure factors of all complexes have been deposited to the protein data bank under accession codes 6QAS, 6QAT, 6QAU and 6QAV.

## ACKNOWLEDGMENT

AC and SK are grateful for support by the SGC, a registered charity (number 1097737) that receives funds from AbbVie, Bayer Pharma AG, Boehringer Ingelheim, Canada Foundation for Innovation, Eshelman Institute for Innovation, Genome Canada, Innovative Medicines Initiative (EU/EFPIA) [ULTRA-DD grant no. 115766], Janssen, Merck KGaA, Germany, MSD, Novartis Pharma AG, Ontario Ministry of Economic Development and Innovation, Pfizer, São Paulo Research Foundation-FAPESP, Takeda, Wellcome [106169/ZZ14/Z]. We are grateful to the DFG funded “center of excellence” (CEF, to SK), the Collaborative Sonderforschungsbereich 1177 Autophagy (SFB1177, to AC, CHB and SK) at Frankfurt University, as well as the German Cancer Consortium (DKTK to CHB and SK), the Deutsche José Carreras Leukämie-Stiftung (DJCLS R14/14, to SS and CHB) and the Else Kröner-Forschungskolleg (to SS). The authors thank staffs at BESSY II and SLS for their support during crystallographic X-ray diffraction test and data collection. The data collection at SLS has been supported by the funding from the European Union’s Horizon 2020 research and innovation program under grant agreement number 730872, project CALIPSOplus.

## ABBREVIATIONS

ULK1: unc-51-like autophagy activating kinase 1
ULK2: unc-51-like autophagy activating kinase 2
GFP: green fluorescent protein
RFP: red fluorescent protein
AML: acute myeloid leukemia
SLK: STE20-like serine/threonine-protein kinase
LOK: Lymphocyte-oriented kinase
DAPK3: Death-associated protein kinase 3
CHK2: Checkpoint kinase 2
ATG1: autophagy-related protein 1
ATG13: autophagy-related protein 13
RB1CC1: RB (Retinoblastoma) 1-inducible coiled-coil protein 1

