## Supplementary information for "Conservation of structure, function and inhibitor binding in UNC-51-like kinase 1 and 2 (ULK1/2)"

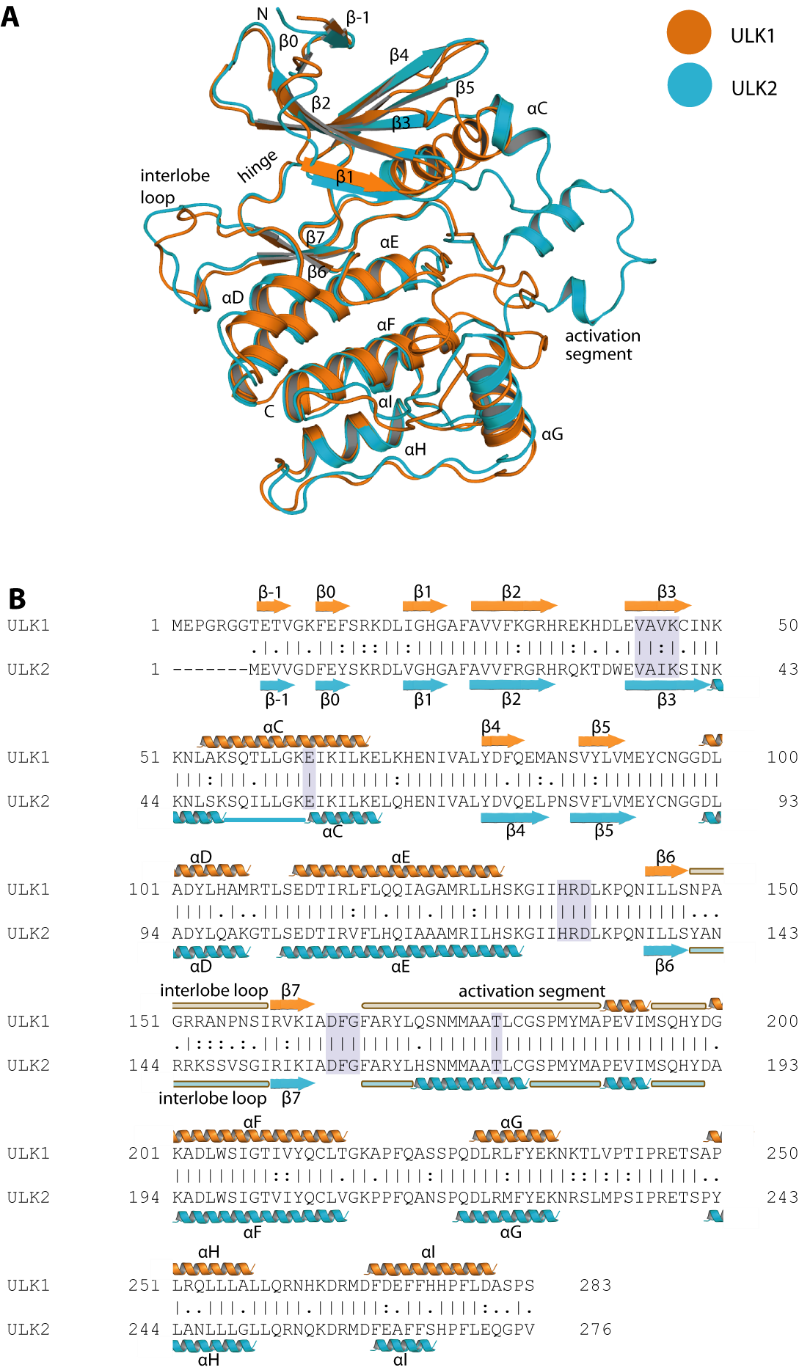

**Supplementary figure 1.** Structure and sequence comparison between ULK1 and ULK2. A) Superimposition of ULK1 and ULK2 kinase domains. B) Sequence alignment with secondary annotations. The purple boxes highlights key catalytic features within the kinases.

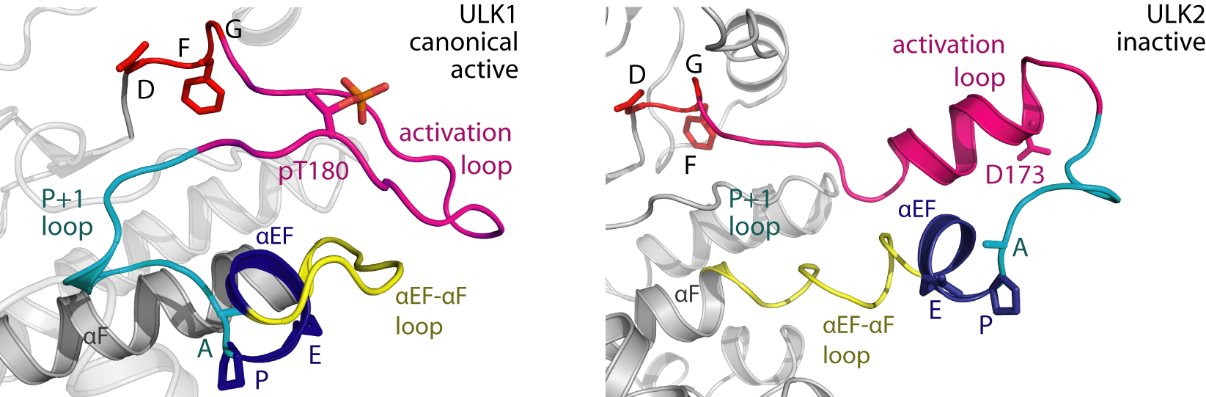

**Supplementary figure 2.** Activation loop conformation in active ULK1 and inactive, dimeric ULK2

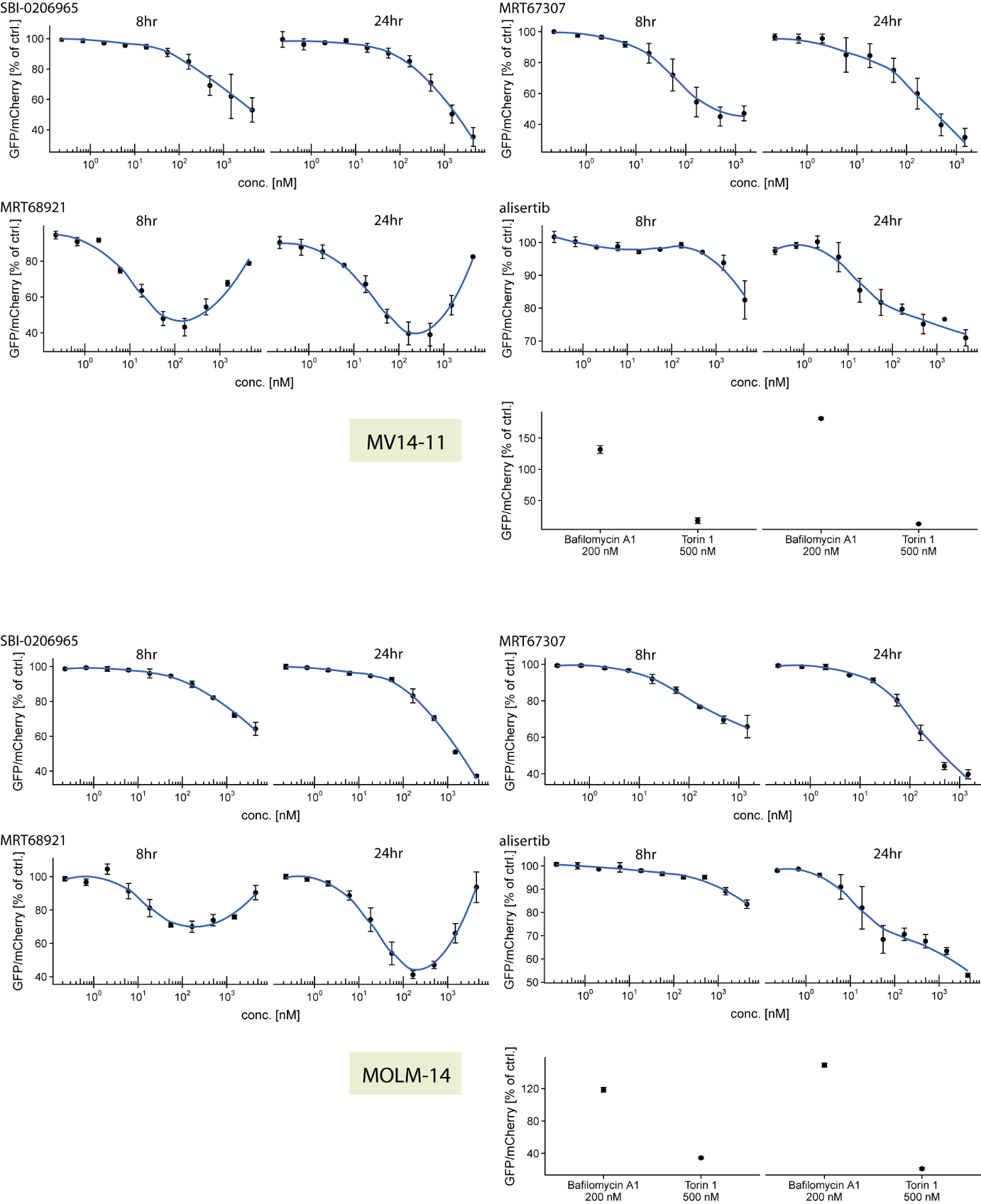

**Supplementary figure 3.** Autophagic flux upon inhibitor treatment in leukemia MV14-11 (top half) and MOLM-14 (bottom half) cell lines measured at two time points. Torin 1 and bafilomycin are used as reference for autophagy induction and inhibition, respectively.

**Supplementary table 1.** Tm shift values for ULK1, ULK2 and Aurora A against diverse 384 kinase inhibitors.

| ULK1 | ULK2 | Aurora A | compound | Targeted kinases | vendor | CatNo. | SMILES |
| --- | --- | --- | --- | --- | --- | --- | --- |
| 12.07 | 14.8 | 6.84 | MRT68921 | ULK1/2 | MedChemExpress | HY-100006A | [H]Cl.O=C(C1CCC1)NCCCNC2=NC(NC3=CC4=C(CN(C)CC4)C=C3)=NC=C2C5CC5.[H]Cl |
| 9.93 | 7.54 | 8.72 | AP26113 | ALK | SelleckChem | S7000 | C1(N=C(C(=CN=1)Cl)NC2C(=CC=CC=2)P(=O)(C)C)NC3C(=CC(=CC=3)N4CCC(CC4)N(C)C)OC |
| 9.75 | 6.49 | 13.52 | SBI-0206965 | ULK1 | SelleckChem | S7885 | CNC(C1=CC=CC=C1OC2=NC(NC3=CC(OC)=C(OC)C(OC)=C3)=NC=C2Br)=O |
| 9.63 | 7.76 | 15.23 | Hesperadin | Aurora Kinase | SelleckChem | S1529 | C1(=CC=C2C(=C1)/C(C(N2)=O)=C(\C3=CC=CC=C3)NC4=CC=C(C=C4)CN5CCCCC5)NS(CC)(=O)=O |
| 9.56 | 9.45 | 16.94 | PF-03814735 | Aurora Kinase | SelleckChem | S2725 | O=C(CNC(C)=O)N1[C@@H]2C3=C(C=C(NC4=NC(NC5CCC5)=C(C(F)(F)F)C=N4)C=C3)[C@H]1CC2 |
| 9.2 | 6.68 | 5.12 | Dabrafenib (GSK2118436) | Raf | SelleckChem | S2807 | C1(C(=CC=CC=1F)F)S(=O)(=O)NC2C(=C(C=CC=2)C3N=C(SC=3C4=NC(=NC=C4)N)C(C)(C)C)F |
| 8.63 | 7.01 | 14.51 | PHA-680632 | Aurora Kinase | SelleckChem | S1454 | C1(=CC=C(C=C1)C(=O)NC2C3=C(NN=2)CN(C3)C(=O)NC4=C(C=CC=C4CC)CC)N5CCN(CC5)C |
| 8.35 | 7.73 | 13.1 | GW8510 | CDK2 | Sigma Aldrich | G7791 | C1=CC(=C(C(=C1N1)C(C1=O)=CNC(=CC=C(C1)S(NC(=CC=CC2)N=2)(=O)=O)C=1)S1)N=C1 |
| 8.35 | 5.64 | 13.62 | TAE684 (NVP-TAE684) | ALK | SelleckChem | S1108 | C1=CC(=CC(=C1NC2=NC=C(C(=N2)NC3=C(C=CC=C3)S(=O)(=O)C(C)C)Cl)OC)N4CCC(CC4)N5CCN(CC5)C |
| 8.11 | 8.13 | 9.2 | MRT67307 | ULK1/2, IKKε and TBK-1 | MedChemExpress | HY-13018 | O=C(NCCCNC1=NC(NC2=CC=CC(CN3CCOCC3)=C2)=NC=C1C4CC4)C5CCC5 |
| 7.93 | 6.41 | 9.7 | TG101209 | JAK,FLT3,c-RET | SelleckChem | S2692 | C1=C(C=C(C=C1)NC2=NC(=NC=C2C)NC3=CC=C(C=C3)N4CCN(CC4)C)S(NC(C)(C)C)(=O)=O |
| 7.68 | 4.16 | 14.91 | TAE226 (NVP-TAE226) | FAK | SelleckChem | S2820 | C1(=CC(=C(C=C1)NC2=NC(=C(C=N2)Cl)NC3=C(C=CC=C3)C(NC)=O)OC)N4CCOCC4 |
| 7.66 | 6.58 | 12.47 | OTSSP167 | MELK | MedChemExpress | HY-15512A | CC(c1cnc2ccc(c3cc(c(c(c3)[Cl])O)[Cl])nc2c1N[C@@H]1CC[C@H](CC1)CN(C)C)=O |
| 7.55 | 7.7 | 5.76 | Sunitinib malate | VEGFR2, PDGFRβ (RTK) | SelleckChem | S1042 | CCN(CC)CCNC(c1c(COC(C[C@@H](C(O)=O)O)=O)c(C=C2C(Nc3ccc(cc23)F)=O)[nH]c1C)=O |
| 7.28 | 4.87 | 14.28 | BX-795 | IκB/IKK,PDK-1 | SelleckChem | S1274 | N1(CCCC1)C(=O)NC2C=C(C=CC=2)NC3N=C(C(=CN=3)I)NCCCNC(C4=CC=CS4)=O |
| 7.09 | 5.56 | 6.18 | GZD824 | Bcr-Abl | SelleckChem | S7194 | C1(=CC(=CC(=C1C)C#CC2=CC3=C(N=C2)NN=C3)C(NC4=CC(=C(C=C4)CN5CCN(CC5)C)C(F)(F)F)=O).CS(=O)(=O)O.CS(=O)(=O)O |
| 6.46 | 5.66 | 7.96 | AZD7762 | Chk | SelleckChem | S1532 | C1(=CC=CC(=C1)C2SC(=C(C=2)NC(N)=O)C(N[C@@H]3CNCCC3)=O)F |
| 6.42 | 5.54 | 7.35 | AZD3463 | ALK | SelleckChem | S7106 | C12=C(C=CC=C1)C(=CN2)C3=C(C=NC(=N3)NC4=C(C=C(C=C4)N5CCC(CC5)N)OC)Cl |
| 6.31 | 2.99 | 1.14 | MLN0905 | PLK1 | SelleckChem | S2898 | CC1=NC=C(CCCN(C)C)C=C1NC2=NC(C(C=CC(C(F)(F)F)=C3)=C3NC(C4)=S)=C4C=N2 |
| 6.17 | 3.26 | 11.63 | R406 (free base) | Syk | SelleckChem | S1533 | C1(=C(C=C(C=C1OC)NC2=NC(=C(C=N2)F)NC3=NC4=C(C=C3)OC(C(N4)=O)(C)C)OC)OC |
| 5.92 | 5.89 | 4.99 | CHIR-124 | Chk | SelleckChem | S2683 | C1(C(=C(C2=C(N1)C=CC(=C2)Cl)N[C@H]3C4CCN(C3)CC4)C5=NC6=C(N5)C=CC=C6)=O |
| 5.6 | 1.77 | 3.95 | BMS-794833 | VEGFR,c-Met | SelleckChem | S2201 | N1=C(C(=C(C=C1)OC2=CC=C(C=C2F)NC(C3=CNC=C(C3=O)C4=CC=C(C=C4)F)=O)Cl)N |
| 5.34 | 5.12 | 3.99 | Crenolanib (CP-868596) | PDGFR | SelleckChem | S2730 | C1=CC(=C2C(=C1)C=CC(=N2)N3C=NC4=C3C=CC(=C4)OCC5(COC5)C)N6CCC(CC6)N |
| 5.33 | 6.6 | 14.29 | XMU-MP-1 | MST1/2 | MedChemExpress | HY-100526 | O=S(C1=CC=C(NC2=NC=C(C(N(C)C3=C4SC=C3)=N2)N(C)C4=O)C=C1)(N)=O |
| 5.3 | 4.18 | 6.57 | Milciclib (PHA-848125) | CDK | SelleckChem | S2751 | C12CC(C3=C(C(=NC(=NC=1)NC4=CC=C(C=C4)N5CCN(CC5)C)2)N(N=C3C(NC)=O)C)(C)C |
| 5.21 | 4.71 | 8.82 | Cdk1/2 Inhibitor III | CDK1/2 | Calbiochem (EMD) | 217714 | O=S(=O)(N)C(C=CC1NC(=NN(C2N)C(=S)NC(C(=CC3)F)=C(C=3)F)N=2)=CC=1 |
| 4.97 | 2.26 | 14.1 | CYC116 | Aurora Kinase,VEGFR | SelleckChem | S1171 | C1(=NC=CC(=N1)C2SC(=NC=2C)N)NC3=CC=C(C=C3)N4CCOCC4 |
| 4.83 | 3.74 | 14.86 | AT9283 | JAK,Aurora Kinase,Bcr-Abl | SelleckChem | S1134 | N(C(=O)NC1C(=NNC=1)C2NC3C(N=2)=CC(=CC=3)CN4CCOCC4)C5CC5 |
| 4.64 | 5.34 | 4.52 | Purvalanol A | CDK | SelleckChem | S7793 | N(C(=C(C(=NC1NC(CO)C(C)C)N2C(C)C)N=C2)N=1)C(=CC(=CC1)[Cl])C=1 |
| 4.58 | 4.53 | 12.15 | PF-573228 | FAK | SelleckChem | S2013 | N1C(CCC2C=C(C=CC1=2)NC3N=C(C(=CN=3)C(F)(F)F)NCC4C=C(C=CC=4)S(=O)(=O)C)=O |
| 4.53 | 2.36 | 8.45 | R406 | Syk,FLT3 | SelleckChem | S2194 | C1(=C(C=C(C=C1OC)NC2=NC(=C(C=N2)F)NC3=NC4=C(C=C3)OC(C(N4)=O)(C)C)OC)OC.C5=CC=CC(=C5)S(O)(=O)=O |
| 4.4 | 4.3 | 4.09 | CP-673451 | PDGFR | SelleckChem | S1536 | C1=CC(=C2C(=C1)C=CC(=N2)N3C4=CC=C(C=C4N=C3)OCCOC)N5CCC(CC5)N |
| 4.32 | 4.85 | 5.53 | Dovitinib (TKI-258) Dilactic Acid | PDGFR,FGFR,c-Kit,FLT3,VEGFR | SelleckChem | S2769 | N1(C(C(=C(C2C(=CC=CC1=2)F)N)C3=NC4C(N3)=CC(=CC=4)N5CCN(CC5)C)=O).CC(C(O)=O)O.CC(C(O)=O)O |
| 4.18 | 4.6 | 7.34 | BI-D1870 | S6 Kinase | SelleckChem | S2843 | C1(=C(C=C(C=C1F)NC2=NC3=C(C=N2)N(C(C(N3CCC(C)C)C)=O)C)F)O |
| 4.02 | 3.63 | 7.66 | TG101348 (SAR302503) | JAK | SelleckChem | S2736 | C1(=CC=C(C=C1)NC2=NC(=C(C=N2)C)NC3=CC=CC(=C3)S(NC(C)(C)C)(=O)=O)OCCN4CCCC4 |
| 3.95 | 3.3 | 9.18 | KW-2449 | Aurora Kinase,Bcr-Abl,FLT3 | SelleckChem | S2158 | C1=CC=CC2=C1NN=C2/C=C/C3=CC=C(C=C3)C(=O)N4CCNCC4 |
| 3.95 | 2.74 | 5.01 | CEP-33779 | JAK | SelleckChem | S2806 | C1(=CC=CC(=C1)NC2N=C3N(N=2)C=CC=C3C4=CC=C(C=C4)S(=O)(C)=O)N5CCN(CC5)C |
| 3.91 | 2.87 | 2.85 | 5-Iodotubercidin; 4-Amino-5-iodo-7-(b-D-ribofuranosyl)pyrrolo[2,3-d]pyrimidine | haspin, adenosine kinase | Calbiochem (EMD) | 407900 | C(=NC=N1)(C(=C1N(C1)C(C(C(C2CO)O)O)O2)C=1I)N |
| 3.83 | 0.88 | 14.1 | AMG 900 | Aurora Kinase | SelleckChem | S2719 | C1=CC(=C(N=C1)OC2=CC=C(C=C2)NC3C4=C(C(=NN=3)C5SC=C(C=5)C)C=CC=C4)C6=NC(=NC=C6)N |
| 3.53 | 1.37 | 3.75 | LDK378 | ALK | SelleckChem | S7083 | C1=CC(=C(C=C1)NC2=C(Cl)C=NC(=N2)NC3=C(C=C(C(=C3)C)C4CCNCC4)OC(C)C)S(C(C)C)(=O)=O |
| 3.49 | 1.86 | 14.59 | BX-912 | PDK-1 | SelleckChem | S1275 | N1(CCCC1)C(=O)NC2C=C(C=CC=2)NC3N=C(C(=CN=3)Br)NCCC4=CN=CN4 |
| 3.32 | 5.02 | 8.19 | BMS-536924 | IGF-1R/IR | SelleckChem | S1012 | Cc1cc(cc2c1nc(C1=C(C=CNC1=O)NC[C@H](c1cccc(c1)[Cl])O)[nH]2)N1CCOCC1 |
| 3.32 | 3.21 | 17.45 | Danusertib (PHA-739358) | c-RET,FGFR,Bcr-Abl,Aurora Kinase | SelleckChem | S1107 | C1(=CC=CC=C1)[C@H](C(N2CC3=C(C2)C(=NN3)NC(=O)C4=CC=C(C=C4)N5CCN(CC5)C)=O)OC |
| 3.26 | 2.98 | 4.05 | NVP-BSK805 2HCl | JAK | SelleckChem | S2686 | C1(=CC=C2C(=C1C3=CC(=C(C(=C3)F)CN4CCOCC4)F)N=C(C=N2)C5=CN(N=C5)C6CCNCC6).Cl.Cl |
| 3.23 | 2.99 | 10.35 | SU11274 | c-Met | SelleckChem | S1080 | C1(=CC=CC(=C1)N(C)S(C2=CC3=C(C=C2)NC(C/3=C/C4NC(=C(C=4C)C(N5CCN(CC5)C)=O)C)=O)(=O)=O)Cl |
| 3.06 | 2.02 | 13.42 | JNJ-7706621 | CDK,Aurora Kinase | SelleckChem | S1249 | C1(=CC=C(C=C1)NC2N=C(N(N=2)C(C3=C(C=CC=C3F)F)=O)N)S(N)(=O)=O |
| 2.97 | 1.64 | 6.99 | FM-381 | JAK3 | MedChemExpress | HY-102046 | CN(C)C(C(=Cc1ccc(c2nc3cnc4c(cc[nH]4)c3n2C2CCCCC2)o1)C#N)=O |
| 2.83 | 1.57 | 1.2 | Sorafenib | Raf-1, B-Raf and VEGFR-2 | SelleckChem | S7397 | C(=C(C(=CC1)[Cl])C(F)(F)F)C=1NC(NC(=CC=C(C1)OC(=CC(=NC2)C(NC)=O)C=2)C=1)=O |
| 2.8 | 0.82 | 7.42 | BMS-777607 | Axl,c-Met | SelleckChem | S1561 | C1(=C(N=CC=C1OC2=CC=C(C=C2F)NC(=O)C3C(N(C=CC=3OCC)C4=CC=C(C=C4)F)=O)N)Cl |
| 2.73 | 1.27 | 9.92 | XL019 | JAK | SelleckChem | S7036 | C1=CC(=CC=C1NC([C@@H]2CCCN2)=O)C3=CC=NC(=N3)NC4=CC=C(C=C4)N5CCOCC5 |
| 2.59 | -0.5 | 2.21 | Tyrphostin AG 879 | HER2 | SelleckChem | S2816 | C(/C(=C/C1C=C(C(=C(C=1)C(C)(C)C)O)C(C)(C)C)C#N)(N)=S |
| 2.49 | 1.82 | 12.48 | WZ3146 | EGFR | SelleckChem | S1170 | C1(=CC=C(C=C1)NC2=NC=C(C(=N2)OC3=CC(=CC=C3)NC(C=C)=O)Cl)N4CCN(CC4)C |
| 2.46 | 0.75 | 9.95 | Crizotinib (PF-02341066) | c-Met,ALK | SelleckChem | S1068 | C1(=CC(=C(N=C1)N)O[C@@H](C2=C(C=CC(=C2Cl)F)Cl)C)C3=CN(N=C3)C4CCNCC4 |
| 2.45 | 2.06 | 10.76 | PF-00562271 | FAK | SelleckChem | S2672 | C1(=NC(=C(C=C1)CNC2=C(C=NC(=N2)NC3=CC=C4C(=C3)CC(N4)=O)C(F)(F)F)N(C)S(=O)(C)=O).C5=CC=CC(=C5)S(O)(=O)=O |
| 2.43 | 2.24 | 11.08 | PF-562271 | FAK | SelleckChem | S2890 | C1=NC(=C(C=C1)CNC2=C(C=NC(=N2)NC3=CC=C4C(=C3)CC(N4)=O)C(F)(F)F)N(C)S(=O)(=O)C |
| 2.37 | 0.96 | 9.56 | IKK-16 (IKK Inhibitor VII) | IκB/IKK | SelleckChem | S2882 | C1(C=CC(=CC=1)NC2N=C(C=CN=2)C3=CC4C(S3)=CC=CC=4)C(=O)N5CCC(CC5)N6CCCC6 |
| 2.34 | 1.22 | 4.98 | LY2835219 | CDK | SelleckChem | S7158 | C1(=CC=C(N=C1)NC2=NC(=C(C=N2)F)C3=CC(=C4C(=C3)N(C(=N4)C)C(C)C)F)CN5(CCN(CC5)CC).CS(=O)(=O)O |
| 2.29 | 1.32 | 13.73 | BMS-754807 | IGF-1R,Trk receptor,c-Met | SelleckChem | S1124 | C12N(N=C(N=C1NC3C=C(NN=3)C4CC4)N5CCC[C@]5(C)C(=O)NC6=CC=C(N=C6)F)C=CC=2 |
| 2.28 | 1.84 | 11.61 | SNS-314 | Aurora Kinase | SelleckChem | S1154 | C12(=C(N=CN=C1NCCC3SC(=NC=3)NC(NC4=CC(=CC=C4)Cl)=O)C=CS2).CS(=O)(=O)O |
| 2.11 | 1.81 | 15.73 | TAK-901 | Aurora Kinase | SelleckChem | S2718 | C1N(CCC(C1)NC(C2=C(C3=C(C(=C2)C4=CC(=CC=C4)S(CC)(=O)=O)C5=C(N3)N=CC(=C5)C)C)=O)C |
| 2.1 | 2.4 | 11.1 | PF-477736 | Chk | SelleckChem | S2904 | C1=C(C=C2C3=C1C(NN=CC(=C(N2)C4C=NN(C=4)C)3)=O)NC([C@@H](C5CCCCC5)N)=O |
| 2.07 | 1.46 | 0.96 | Bosutinib (SKI-606) | Src | SelleckChem | S1014 | N1=CC(=C(C2=CC(=C(C=C12)OCCCN3CCN(CC3)C)OC)NC4C(=CC(=C(C=4)OC)Cl)Cl)C#N |
| 1.97 | -3.86 | 3.13 | Tyrphostin 9 | EGFR | SelleckChem | S2895 | C1(C(=C(C=C(C=1)C=C(C#N)C#N)C(C)(C)C)O)C(C)(C)C |
| 1.93 | 0.16 | -0.59 | TX-1123; 2-((3,5-di-tert-Butyl-4-hydroxyphenyl)-methylene)-4-cyclopentene-1,3-dione | c-Src, eEF2-K, and PKA | Calbiochem (EMD) | 655200 | C(=C(C(=C(C1)C(C)(C)C)O)C(C)(C)C)C=1C=C(C(C=C1)=O)C1=O |
| 1.87 | 1.41 | 4.12 | Vemurafenib (PLX4032, RG7204) | Raf | SelleckChem | S1267 | C1(=CN=C2C(=C1)C(=CN2)C(=O)C3=C(C(=CC=C3F)NS(=O)(=O)CCC)F)C4=CC=C(C=C4)Cl |
| 1.74 | 0.24 | 6.36 | CYT387 | JAK | SelleckChem | S2219 | C1=C(C=CC(=C1)NC2=NC=CC(=N2)C3=CC=C(C=C3)C(NCC#N)=O)N4CCOCC4 |
| 1.67 | 1.21 | 9.56 | AZD5438 | CDK | SelleckChem | S2621 | C1=C(C=CC(=C1)NC2=NC=CC(=N2)C3=CN=C(N3C(C)C)C)S(=O)(C)=O |
| 1.67 | 1.04 | 12.46 | CCT137690 | Aurora Kinase | SelleckChem | S2744 | C1(=CN=C2C(=C1N3CCN(CC3)CC4C=C(ON=4)C)N=C(N2)C5=CC=C(C=C5)N6CCN(CC6)C)Br |
| 1.52 | 2.3 | 13.96 | ENMD-2076 | Aurora Kinase,FLT3,VEGFR | SelleckChem | S1181 | C1(N=C(C=C(N=1)N2CCN(CC2)C)NC3=NNC(=C3)C)/C=C/C4C=CC=CC=4 |
| 1.5 | 0.51 | 4.93 | BMS-265246 | CDK | SelleckChem | S2014 | C(=O)(C1C(=CC(=CC=1F)C)F)C2C(=C3C(=NC=2)NN=C3)OCCCC |
| 1.45 | 0.32 | 1.53 | Quercetin; 3,3',4',5,7-Pentahydroxyflavone | promiscuous | Calbiochem (EMD) | 551600 | C1=C(C=C(C(=C1O1)C(C(=C1C(=CC=C(C1O)O)C=1)O)=O)O)O |
| 1.45 | 0.92 | 12.29 | GSK1070916 | Aurora Kinase | SelleckChem | S2740 | C1=CN=C2C(=C1C3=CN(N=C3C4=CC=C(C=C4)NC(N(C)C)=O)CC)C=C(N2)C5=CC=CC(=C5)CN(C)C |
| 1.44 | -0.17 | 1.11 | 3-(2-Chloro-3-indolylmethylene)-1,3-dihydroindol-2-one | CDK1 | Calbiochem (EMD) | 217695 | C1=CC=CC(=C1C(=C1[Cl])C=C(C(N2)=O)C(=C2C=CC2)C=2)N1 |
| 1.41 | 0.96 | 1.01 | DCC-2036 (Rebastinib) | Bcr-Abl | SelleckChem | S2634 | C1(=C(C=C(C=C1)OC2=CC(=NC=C2)C(NC)=O)F)NC(NC3N(N=C(C=3)C(C)(C)C)C4=CC5=C(C=C4)N=CC=C5)=O |
| 1.32 | 0.65 | 1.19 | SD 208 | TGF-βRI | SelleckChem | S7624 | N1=CC=NC(=C1C1NC(=CC=NC2)C=2)N=C(N=1)C(=CC(=CC1)[Cl])C=1F |
| 1.25 | 0.39 | 17.18 | LY2784544 | JAK | SelleckChem | S2179 | C1C(=NN2C(C=1CN3CCOCC3)=NC(=C2CC4=CC=C(C=C4F)Cl)C)NC5NN=C(C=5)C |
| 1.24 | 0.47 | 0.66 | Dinaciclib (SCH727965) | CDK | SelleckChem | S2768 | C1C[C@H](N(CC1)C2C=C(N3C(N=2)=C(C=N3)CC)NCC4=CC=C[N+](=C4)[O-])CCO |
| 1.21 | 0.23 | 2.79 | Sorafenib Tosylate | PDGFR,Raf,VEGFR | SelleckChem | S1040 | C1(=CC=C(C=C1C(F)(F)F)NC(NC2=CC=C(C=C2)OC3=CC=NC(=C3)C(NC)=O)=O)Cl.C4=CC(=CC=C4S(O)(=O)=O)C |
| 1.2 | 0.42 | 14.74 | AZ 960 | JAK | SelleckChem | S2214 | C1=C(C(=NC(=C1F)NC2C=C(NN=2)C)N[C@H](C3=CC=C(C=C3)F)C)C#N |
| 1.18 | 0.28 | 4.55 | Ruxolitinib (INCB018424) | JAK | SelleckChem | S1378 | N1=CN=C2C(=C1C3=CN(N=C3)[C@@H](CC#N)C4CCCC4)C=CN2 |
| 1.16 | 0.95 | 0.61 | Cdc7/CDK9 inhibitor | CDK7/9 | Calbiochem (EMD) | 217707 | C(C(=C(C(N1)=O)C2)NC=2C(=CC=NC2)C=2)C1 |
| 1.16 | 1.57 | 4.09 | SCH772984 | ERK1/2 | Small Molecules | 33-2262 | C1CN(CC(N2CCN(CC2)c2ccc(cc2)c2ncccn2)=O)C[C@@H]1C(Nc1ccc2c(c1)c(c1ccncc1)n[nH]2)=O |
| 1.13 | 2.13 | 0.88 | Palbociclib (PD-0332991) HCl | CDK | SelleckChem | S1116 | N1(=C(N=C2C(=C1)C(=C(C(N2C3CCCC3)=O)C(C)=O)C)NC4=NC=C(C=C4)N5CCNCC5).Cl |
| 1.11 | 1.58 | 2.43 | MK-8776 (SCH 900776) | CDK,Chk | SelleckChem | S2735 | C12N=C(C(=C(N1N=CC=2C3=CN(N=C3)C)N)Br)[C@H]4CNCCC4 |
| 1.07 | 1.1 | 6.46 | Foretinib (GSK1363089) | VEGFR,c-Met | SelleckChem | S1111 | C1(=CC=C(C=C1)F)NC(C2(C(NC3=CC(=C(C=C3)OC4C5=C(N=CC=4)C=C(C(=C5)OC)OCCCN6CCOCC6)F)=O)CC2)=O |
| 1.07 | 0.64 | 1.56 | AG-1024 (Tyrphostin) | IGF-1R | SelleckChem | S1234 | C(C(C#N)=CC1C=C(C(=C(C=1)C(C)(C)C)O)Br)#N |
| 1.04 | 1.18 | 11.81 | Fostamatinib (R788) | Syk | SelleckChem | S2625 | C1(=C(C(=CC(=C1)NC2=NC=C(C(=N2)NC3=CC=C4C(=N3)N(C(C(O4)(C)C)=O)COP(O)(O)=O)F)OC)OC)OC |
| 1.01 | 0.08 | 4.44 | S-Ruxolitinib (INCB018424) | JAK | SelleckChem | S2902 | N1C2=C(C(=NC=1)C3=CN(N=C3)[C@H](C4CCCC4)CC#N)C=CN2 |
| 1.01 | 1.46 | 1.49 | PRT062607 (P505-15, BIIB057) HCl | Syk | SelleckChem | S8032 | N1(=C(N=C(C(=C1)C(N)=O)NC2=CC(=CC=C2)N3N=CC=N3)N[C@@H]4CCCC[C@H]4N).Cl |
| 0.93 | 0.41 | 2.04 | H-89; N-[2-((p-Bromocinnamyl)amino)ethyl]-5-isoquinolinesulfonamide | PKA | Calbiochem (EMD) | 371963 | C1=NC=CC(=C1C1)C(=CC=1)S(NCCNCC=CC(=CC=C(C1)[Br])C=1)(=O)=O.[Cl].[Cl] |
| 0.91 | 1.36 | 0.9 | PHA-793887 | CDK | SelleckChem | S1487 | N1N=C(C2=C1C(N(C2)C(C3CCN(CC3)C)=O)(C)C)NC(CC(C)C)=O |
| 0.8 | 0.68 | 4.93 | PLX-4720 | Raf | SelleckChem | S1152 | C1(=CN=C2C(=C1)C(=CN2)C(=O)C3=C(C(=CC=C3F)NS(=O)(=O)CCC)F)Cl |
| 0.77 | 0.46 | 14.07 | WZ8040 | EGFR | SelleckChem | S1179 | C1(=CC=C(C=C1)NC2=NC=C(C(=N2)SC3=CC(=CC=C3)NC(C=C)=O)Cl)N4CCN(CC4)C |
| 0.71 | 0.1 | 4.8 | MK-2461 | c-Met,PDGFR,FGFR | SelleckChem | S2774 | C12=C(C(C3=C(C=C1)C=CC(=C3)NS(N(C[C@H]4OCCOC4)C)(=O)=O)=O)C=C(C=N2)C5=CN(N=C5)C |
| 0.6 | 1.24 | 18.79 | VX-680 (MK-0457, Tozasertib) | Aurora Kinase | SelleckChem | S1048 | C1(=CC(=NC(=N1)SC2=CC=C(C=C2)NC(=O)C3CC3)N4CCN(CC4)C)NC5C=C(NN=5)C |
| 0.6 | 0.18 | 3.45 | PIK-75 | PI3K,DNA-PK | SelleckChem | S1205 | C1(C(=CN2C(C=1)=NC=C2/C=N/N(S(C3=C(C=CC(=C3)[N+](=O)[O-])C)(=O)=O)C)Br).Cl |
| 0.57 | 0.37 | 3.84 | WZ4003 | AMPK | SelleckChem | S7317 | N1=C(N=C(C(=C1)Cl)OC2=CC(=CC=C2)NC(CC)=O)NC3=C(C=C(C=C3)N4CCN(CC4)C)OC |
| 0.53 | 0.29 | 1.64 | Ponatinib (AP24534) | PDGFR,FGFR,VEGFR,Bcr-Abl | SelleckChem | S1490 | C1C=NN2C(C=1)=NC=C2C#CC3=C(C=CC(=C3)C(=O)NC4=CC(=C(C=C4)CN5CCN(CC5)C)C(F)(F)F)C |
| 0.53 | -0.01 | 7.97 | TWS119 | GSK-3 | SelleckChem | S1590 | C1=CC=C(C=C1O)OC2=NC=NC3=C2C=C(N3)C4=CC=CC(=C4)N |
| 0.49 | -0.06 | 0.95 | PQ401 | IGF-1R | SelleckChem | S8003 | N(C(=O)NC1C=C(N=C2C=CC=CC=12)C)C3=C(C=CC(=C3)Cl)OC |
| 0.46 | 0.22 | 9.65 | Axitinib | c-Kit,VEGFR,PDGFR | SelleckChem | S1005 | C1(=CC=CC=C1C(=O)NC)SC2=CC3=C(C=C2)C(=NN3)/C=C/C4=CC=CC=N4 |
| 0.46 | 0.6 | 3.52 | WZ4002 | EGFR | SelleckChem | S1173 | C1(=CC(=C(C=C1)NC2=NC=C(C(=N2)OC3=CC(=CC=C3)NC(C=C)=O)Cl)OC)N4CCN(CC4)C |
| 0.44 | 0.18 | 0.36 | met-ADP; a,b-Methyleneadenosine 5'diphosphate |  | Sigma Aldrich | M3763 | OC(C(N(C(=C1C(N)=N2)N=C2)C=N1)O1)C(O)C1COP(CP(=O)(O)O)(=O)O |
| 0.42 | -0.09 | 5.07 | AEE788 (NVP-AEE788) | HER2,VEGFR,EGFR | SelleckChem | S1486 | C1=NC(=C2C(=N1)NC(=C2)C3=CC=C(C=C3)CN4CCN(CC4)CC)N[C@@H](C5=CC=CC=C5)C |
| 0.37 | 0.46 | -0.15 | BIRB 796 (Doramapimod) | p38 MAPK | SelleckChem | S1574 | C1=C(C2=C(C(=C1)OCCN3CCOCC3)C=CC=C2)NC(NC4=CC(=NN4C5=CC=C(C=C5)C)C(C)(C)C)=O |
| 0.35 | 0.13 | 0.21 | GDC-0879 | Raf | SelleckChem | S1104 | C(CN1N=C(C(=C1)C2=CC=C3C(=C2)CCC/3=N/O)C4C=CN=CC=4)O |
| 0.35 | 0.65 | 5.83 | TPCA-1 | IκB/IKK | SelleckChem | S2824 | C1(=C(C=C(S1)C2C=CC(=CC=2)F)C(=O)N)NC(=O)N |
| 0.34 | 0.01 | 0.23 | Harmine | DYRK1A | Sigma Aldrich | 51400 | C1C(=CC=C(C=1N1)C(=C1C1C)C=CN=1)OC |
| 0.27 | 0.22 | 4.33 | Dasatinib | Bcr-Abl,c-Kit,Src | SelleckChem | S1021 | N1=C(N=C(C=C1NC2=NC=C(S2)C(NC3=C(C=CC=C3C)Cl)=O)N4CCN(CC4)CCO)C |
| 0.27 | 0.23 | 0.45 | NVP-AEW541 | IGF-1R | SelleckChem | S1034 | C1=CC(=CC(=C1)OCC2=CC=CC=C2)C3=CN(C4=C3C(=NC=N4)N)[C@@H]5C[C@@H](C5)CN6CCC6 |
| 0.27 | 0.1 | 1.91 | Golvatinib (E7050) | VEGFR,c-Met | SelleckChem | S2859 | C1=C(C=CC(=C1F)NC(C2(C(NC3=CC=C(C=C3)F)=O)CC2)=O)OC4=CC(=NC=C4)NC(N5CCC(CC5)N6CCN(CC6)C)=O |
| 0.23 | -0.39 | 1.71 | Butein | EGFR | SelleckChem | S8036 | C1(=C(C=C(C=C1)O)O)C(/C=C/C2=CC(=C(C=C2)O)O)=O |
| 0.22 | 0.8 | 0.71 | Flavopiridol | CDK1/2/4/6/9 | SelleckChem | S1230 | C(=C(C1)O)(C(=C(C=1O)C(C=C1C(=C(C=CC2)[Cl])C=2)=O)O1)C(C(CN(C1)C)O)C1 |
| 0.19 | 0.93 | 9.67 | Barasertib (AZD1152-HQPA) | Aurora Kinase | SelleckChem | S1147 | C1(=CC=C2C(=C1)N=CN=C2NC3NN=C(C=3)CC(NC4=CC=CC(=C4)F)=O)OCCCN(CCO)CC |
| 0.16 | -0.85 | 5.81 | TCS ERK 11e | ERK1/2 | Tocris | 4465 | Cc1cnc(Nc2ccc(cc2[Cl])F)nc1c1cc(C(N[C@H](CO)c2cccc(c2)[Cl])=O)[nH]c1 |
| 0.15 | -0.22 | 23.55 | Alisertib (MLN8237) | Aurora Kinase | SelleckChem | S1133 | C1=CC=C(C(=C1OC)C2=NCC3=C(C4=C2C=C(C=C4)Cl)N=C(N=C3)NC5=CC(=C(C=C5)C(O)=O)OC)F |
| 0.15 | -0.05 | 1.94 | Ro 31-8220 Mesylate | PKC | SelleckChem | S7207 | C1(=C2C(=CC=C1)C(=CN2CCCSC(=N)N)C3C(=O)NC(C(C4=CN(C5=CC=CC=C54)C)=3)=O).CS(=O)(=O)O |
| 0.11 | 0.04 | 0.02 | PNU112455A; Cdk2/5 Inhibitor; N4-(6-Aminopyrimidin-4-yl)-sulfanilamide, HCI | CDK2/5 | Calbiochem (EMD) | 219448 | C(=NC=NC1NC(=CC=C(C2)S(N)(=O)=O)C=2)(C=1)N.[Cl] |
| 0.11 | 0.18 | 0.29 | BS-181 HCl | CDK | SelleckChem | S1572 | C1(=CC(=NC2N1N=CC=2C(C)C)NCCCCCCN)NCC3(=CC=CC=C3).Cl |
| 0.08 | 0.19 | 0.01 | Tubercidin |  | Sigma Aldrich | T0642 | C([C@@H]1[C@H]([C@H]([C@H](n2ccc3c(N)ncnc23)O1)O)O)O |
| 0.05 | 0.35 | 0.34 | AT7519 | CDK | SelleckChem | S1524 | C1=C(C(=C(C=C1)Cl)C(NC2=CNN=C2C(=O)NC3CCNCC3)=O)Cl |
| 0.03 | -0.1 | 0.55 | SB590885 | Raf | SelleckChem | S2220 | N1C(=C(NC=1C2=CC=C(C=C2)OCCN(C)C)C3=CC=C4C(=C3)CCC/4=N/O)C5=CC=NC=C5 |
| 0.03 | 0.44 | 6.84 | Pacritinib (SB1518) | JAK | SelleckChem | S8057 | C1=CN=C2N=C1C3=CC(=CC=C3)COC/C=C/COCC4=CC(=CC=C4OCCN5CCCC5)N2 |
| 0.01 | 0.24 | 4.48 | AZD4547 | FGFR | SelleckChem | S2801 | C1(=CC(=CC(=C1)CCC2=CC(=NN2)NC(=O)C3=CC=C(C=C3)N4C[C@@H](N[C@@H](C4)C)C)OC)OC |
| -0.01 | -0.14 | 1.22 | Pelitinib (EKB-569) | EGFR | SelleckChem | S1392 | C1(=C(C=C2C(=C1)N=CC(=C2NC3=CC(=C(C=C3)F)Cl)C#N)NC(/C=C/CN(C)C)=O)OCC |
| -0.02 | -0.46 | 9.49 | CNX-774 | BTK | SelleckChem | S7257 | C1=C(C=C(C=C1)NC(C=C)=O)NC2=C(C=NC(=N2)NC3=CC=C(C=C3)OC4=CC=NC(=C4)C(NC)=O)F |
| -0.03 | -0.53 | 1.68 | Regorafenib (BAY 73-4506) | c-RET,VEGFR | SelleckChem | S1178 | C1=NC(=CC(=C1)OC2=CC(=C(C=C2)NC(NC3=CC=C(C(=C3)C(F)(F)F)Cl)=O)F)C(=O)NC |
| -0.04 | -0.27 | 3.89 | AZD1080 | GSK-3 | SelleckChem | S7145 | C1=C(C=C2C(=C1)NC(=C2C3=CC=C(C=N3)CN4CCOCC4)O)C#N |
| -0.06 | -0.44 | -0.15 | GW5074 | c-Raf | MedChemExpress | HY-10542 | O=C1NC2=C(/C1=C/C3=CC(Br)=C(C(Br)=C3)O)C=C(C=C2)I |
| -0.06 | 0.36 | 0.03 | BI6727 (Volasertib) | PLK | SelleckChem | S2235 | [C@H]1(C(N(C2=C(N1C(C)C)N=C(N=C2)NC3=CC=C(C=C3OC)C(N[C@@H]4CC[C@H](CC4)N5CCN(CC5)CC6CC6)=O)C)=O)CC |
| -0.09 | -0.22 | 4.14 | PHA-665752 | c-Met | SelleckChem | S1070 | C1=CC=C(C(=C1Cl)CS(C2=CC3=C(C=C2)NC(C/3=C/C4=C(C(=C(N4)C)C(N5[C@H](CCC5)CN6CCCC6)=O)C)=O)(=O)=O)Cl |
| -0.09 | -0.13 | 7.92 | Cabozantinib (XL184, BMS-907351) | FLT3,Tie-2,c-Kit,c-Met,VEGFR,Axl | SelleckChem | S1119 | C12=C(C(=CC=N1)OC3=CC=C(C=C3)NC(C4(C(NC5=CC=C(C=C5)F)=O)CC4)=O)C=C(C(=C2)OC)OC |
| -0.11 | 0.39 | 2.5 | OSI-906 (Linsitinib) | IGF-1R | SelleckChem | S1091 | C1=C(C=C2C(=C1)C=CC(=N2)C3=CC=CC=C3)C4=C5N(C(=N4)[C@H]6C[C@@](C6)(C)O)C=CN=C5N |
| -0.11 | -0.13 | 11.65 | Tivozanib (AV-951) | VEGFR,PDGFR,c-Kit | SelleckChem | S1207 | C1(=CC=C(C(=C1)Cl)NC(=O)NC2C=C(ON=2)C)OC3=CC=NC4=C3C=C(C(=C4)OC)OC |
| -0.14 | 0.03 | 9.2 | AVL-292 | BTK | SelleckChem | S7173 | C1(=CN=C(N=C1NC2=CC(=CC=C2)NC(C=C)=O)NC3=CC=C(C=C3)OCCOC)F |
| -0.15 | -0.39 | 9.42 | TSU-68 (SU6668, Orantinib) | VEGFR,PDGFR,FGFR | SelleckChem | S1470 | C1=CC=C2C(=C1)/C(C(N2)=O)=C/C3NC(=C(C=3C)CCC(O)=O)C |
| -0.17 | 0.14 | 2.82 | CCT128930 | Akt | SelleckChem | S2635 | N1=CN=C2C(=C1N3CCC(CC3)(N)CC4=CC=C(C=C4)Cl)C=CN2 |
| -0.17 | 0.35 | 0.86 | GSK1838705A | IGF-1R,ALK | SelleckChem | S2703 | N1=C(N=C2C(=C1NC3=CC=CC(=C3C(=O)NC)F)C=CN2)NC4=C(C=C5C(=C4)N(CC5)C(=O)CN(C)C)OC |
| -0.18 | -0.81 | 0.76 | BMS-345541 | IκB/IKK | SelleckChem | S8044 | C1(=CC=C2C(=C1)N3C(C(=N2)NCCN)=NC=C3C)C |
| -0.19 | 0.11 | 2.1 | NVP-ADW742 | IGF-1R | SelleckChem | S1088 | C12=C(N=CN=C1N)N(C=C2C3=CC(=CC=C3)OCC4=CC=CC=C4)[C@@H]5C[C@H](C5)CN6CCCC6 |
| -0.19 | -0.3 | 0.81 | Wortmannin | Autophagy,ATM/ATR,PI3K | SelleckChem | S2758 | C1(C2C3[C@]([C@H](O1)COC)(C4=C(C(C=3OC=2)=O)[C@@]5(CCC([C@@](C[C@@H]4OC(=O)C)5C)=O)[H])C)=O |
| -0.2 | -0.15 | 4.07 | CCT129202 | Aurora Kinase | SelleckChem | S1519 | C1(=CN=C2C(=C1N3CCN(CC3)CC(NC4SC=CN=4)=O)N=C(N2)C5=CC=C(C=C5)N(C)C)Cl |
| -0.21 | 0.23 | 0.74 | AZ 628 | Raf | SelleckChem | S2746 | C1(=CC=CC(=C1)C(NC2=CC=C(C(=C2)NC3=CC4=C(C=C3)N=CN(C4=O)C)C)=O)C(C#N)(C)C |
| -0.22 | -0.58 | 1 | Ki8751 | PDGFR,c-Kit,VEGFR | SelleckChem | S1363 | C1=CN=C2C(=C1OC3=CC=C(C(=C3)F)NC(NC4=CC=C(C=C4F)F)=O)C=C(C(=C2)OC)OC |
| -0.26 | -0.26 | 1.24 | Quercetin | PKC,Src,PI3K,Sirtuin | SelleckChem | S2391 | C1(=CC(=C2C(=C1)OC(=C(C2=O)O)C3=CC=C(C(=C3)O)O)O)O |
| -0.27 | 0.33 | 3.73 | Semaxanib (SU5416) | VEGFR | SelleckChem | S2845 | C1=CC=C2C(=C1)/C(C(N2)=O)=C/C3NC(=CC=3C)C |
| -0.27 | -0.2 | 4.72 | CO-1686 (AVL-301) | EGFR | SelleckChem | S7284 | C1(=CN=C(N=C1NC2=CC=CC(=C2)NC(=O)C=C)NC3=C(C=C(C=C3)N4CCN(CC4)C(C)=O)OC)C(F)(F)F |
| -0.27 | -0.55 | 0.75 | KX2-391 | Src | SelleckChem | S2700 | C1=C(C=CC(=C1)C2=CC=C(N=C2)CC(NCC3=CC=CC=C3)=O)OCCN4CCOCC4 |
| -0.27 | -0.4 | 3.38 | AMG-458 | c-Met | SelleckChem | S2747 | N1(N(C(C(=C1C)C(=O)NC2C=CC(=CN=2)OC3C=CN=C4C=C(C=CC=34)OC)=O)C5C=CC=CC=5)CC(C)(C)O |
| -0.28 | -0.07 | 1.53 | NU6027 | CDK | SelleckChem | S7114 | C1CCCC(C1)COC2=NC(=NC(=C2N=O)N)N |
| -0.29 | -0.4 | 2.57 | Lapatinib (GW-572016) Ditosylate | HER2,EGFR | SelleckChem | S1028 | C1(=C(C=C2C(=C1)N=CN=C2NC3=CC=C(C(=C3)Cl)OCC4=CC=CC(=C4)F)C5OC(=CC=5)CNCCS(=O)(C)=O).C6(=C(C=CC(=C6)S(O)(=O)=O)C).C7=C(C=CC(=C7)S(O)(=O)=O)C |
| -0.31 | -0.5 | 3.21 | VE-822 | ATM/ATR | SelleckChem | S7102 | N1=CC(=NC(=C1N)C2=CC(=NO2)C3=CC=C(C=C3)CNC)C4=CC=C(C=C4)S(=O)(=O)C(C)C |
| -0.32 | -0.37 | 3.37 | VE-821 | ATM/ATR | SelleckChem | S8007 | C1(C(=NC=C(N=1)C2C=CC(=CC=2)S(=O)(=O)C)N)C(=O)NC3C=CC=CC=3 |
| -0.33 | -0.19 | 3.47 | AZD2858 | GSK-3 | SelleckChem | S7253 | C1=C(N=C(C(=N1)N)C(=O)NC2=CN=CC=C2)C3=CC=C(C=C3)S(=O)(=O)N4CCN(CC4)C |
| -0.35 | -0.83 | 1.37 | YM201636 | PI3K | SelleckChem | S1219 | C1C(=CC=C(N=1)N)C(=O)NC2C=C(C=CC=2)C3N=C4C(=C(N=3)N5CCOCC5)OC6C(=CC=CN=6)4 |
| -0.36 | -0.26 | 1.14 | Dacomitinib (PF299804, PF299) | EGFR | SelleckChem | S2727 | C(/C=C/C(NC1=C(C=C2C(=C1)C(=NC=N2)NC3=CC(=C(C=C3)F)Cl)OC)=O)N4CCCCC4 |
| -0.36 | -0.63 | 0.99 | IMD 0354 | IκB/IKK | SelleckChem | S2864 | C1=CC(=CC(=C1O)C(NC2=CC(=CC(=C2)C(F)(F)F)C(F)(F)F)=O)Cl |
| -0.38 | -0.16 | 0.93 | ZM 336372; N-[5-(3-Dimethylaminobenzamido)-2-methylphenyl]-4-hydroxybenzamide | c-Raf | Calbiochem (EMD) | 692000 | C(=CC=CC1N(C)C)(C=1)C(NC(=CC=C(C1NC(C(=CC=C(C2)O)C=2)=O)C)C=1)=O |
| -0.38 | 0.12 | 0.06 | Roscovitine (Seliciclib, CYC202) | CDK | SelleckChem | S1153 | N1=C2C(=C(N=C1N[C@@H](CO)CC)NCC3=CC=CC=C3)N=CN2C(C)C |
| -0.38 | 0.12 | 1.08 | PHA-767491 | CDK | SelleckChem | S2742 | C1=NC=CC(=C1)C2NC3=C(C=2)C(NCC3)=O |
| -0.4 | -1.01 | 1.28 | SPHINX31 | SRPK1 | Cayman | 21582 | C1CN(CCN1Cc1ccccn1)c1ccc(cc1NC(c1ccc(c2ccncc2)o1)=O)C(F)(F)F |
| -0.42 | -0.34 | 4.79 | AZD5363 | Akt | SelleckChem | S8019 | N1(CCC(CC1)(C(=O)N[C@@H](CCO)C2C=CC(=CC=2)Cl)N)C3C4=C(N=CN=3)NC=C4 |
| -0.43 | 0.17 | 0.59 | Chrysophanic Acid | mTOR,EGFR | SelleckChem | S2406 | C1=CC(=C2C(=C1)C(C3=C(C2=O)C(=CC(=C3)C)O)=O)O |
| -0.43 | -0.37 | 3.65 | WHI-P154 | JAK,EGFR | SelleckChem | S2867 | N1C=NC(=C2C=C(C(=CC=12)OC)OC)NC3C=C(C(=CC=3)O)Br |
| -0.45 | -0.77 | 2.14 | OSI-930 | c-Kit,CSF-1R,VEGFR | SelleckChem | S1220 | C1=CC=C2C(=C1)C(=CC=N2)CNC3=C(SC=C3)C(NC4=CC=C(C=C4)OC(F)(F)F)=O |
| -0.46 | -0.7 | 1.24 | SAR245409 (XL765) | PI3K,mTOR | SelleckChem | S1523 | C12=CC=CC=C1N=C(C(=N2)NC3=CC(=CC(=C3)OC)OC)NS(C4=CC=C(C=C4)NC(=O)C5=CC=C(C(=C5)OC)C)(=O)=O |
| -0.46 | -0.06 | 1.06 | SB216763 | GSK-3 | SelleckChem | S1075 | C1=CC=CC2=C1N(C=C2C3=C(C(NC3=O)=O)C4=CC=C(C=C4Cl)Cl)C |
| -0.46 | -0.56 | 4.79 | PIK-294 | PI3K | SelleckChem | S2227 | N1=CN=C2C(=C1N)C(=NN2CC3N(C(C4=C(N=3)C=CC=C4C)=O)C5=C(C=CC=C5)C)C6=CC(=CC=C6)O |
| -0.47 | -0.74 | 8.86 | ACK1 Inhibitor AIM-100 | ACK1 | Calbiochem | 104833 | C1C[C@@H](CNc2c3c(c4ccccc4)c(c4ccccc4)oc3ncn2)OC1 |
| -0.47 | -0.91 | 2.7 | Apatinib | VEGFR | SelleckChem | S2221 | S(=O)(O)(C)=O.C1=CN=C(C(=C1)C(NC2=CC=C(C=C2)C3(CCCC3)C#N)=O)NCC4=CC=NC=C4 |
| -0.49 | -0.4 | 1.84 | GSK1059615 | PI3K,mTOR | SelleckChem | S1360 | C1(=CC=C2C(=C1)C(=CC=N2)C3=CC=NC=C3)/C=C4/C(NC(S4)=O)=O |
| -0.49 | -0.43 | 1.51 | AST-1306 | EGFR | SelleckChem | S2185 | C(C=C)(=O)NC1(=CC2=C(N=CN=C(C=C1)2)NC3C=C(C(=CC=3)OCC4C=C(C=CC=4)F)Cl).C5=CC(=CC=C5S(O)(=O)=O)C |
| -0.49 | -0.57 | 0.74 | U0126-EtOH | MEK | SelleckChem | S1102 | C1(=CC=C(C(=C1)SC(/N)=C(\C(=C(\SC2=C(C=CC=C2)N)N)C#N)C#N)N).CCO |
| -0.51 | -0.36 | 1.08 | BGJ398 (NVP-BGJ398) | FGFR | SelleckChem | S2183 | N(C(=O)NC1C(=C(C=C(C=1Cl)OC)OC)Cl)(C)C2N=CN=C(C=2)NC3C=CC(=CC=3)N4CCN(CC4)CC |
| -0.52 | -0.68 | 16.79 | Aurora A Inhibitor I | Aurora Kinase | SelleckChem | S1451 | N1=CC(=C(N=C1NC2=CC=C(C=C2)CC(N3CCN(CC3)CC)=O)NC4=CC=C(C=C4)C(NC5=CC=CC=C5Cl)=O)F |
| -0.53 | -0.32 | 2.44 | A-674563 | PKA,CDK,Akt | SelleckChem | S2670 | C1=CC=CC(=C1)C[C@@H](COC2=CC(=CN=C2)C3=CC4=C(C=C3)NN=C4C)N |
| -0.53 | -0.11 | 1.06 | SB415286 | GSK-3 | SelleckChem | S2729 | C1(C(=C(C(N1)=O)NC2=CC(=C(C=C2)O)Cl)C3=C(C=CC=C3)[N+](=O)[O-])=O |
| -0.53 | -0.12 | 0.95 | RO3280 | PLK1 | SelleckChem | S7248 | COC1=C(NC2=NC(N(C3CCCC3)CC(F)(F)C(N4C)=O)=C4C=N2)C=CC(C(NC5CCN(C)CC5)=O)=C1 |
| -0.55 | -1.01 | 8.04 | Ibrutinib (PCI-32765) | BTK | SelleckChem | S2680 | C1=NC(=C2C(=N1)N(N=C2C3=CC=C(C=C3)OC4=CC=CC=C4)[C@H]5CN(CCC5)C(C=C)=O)N |
| -0.57 | -0.18 | 2.62 | PP242 | mTOR,Autophagy | SelleckChem | S2218 | N1=CN=C2C(=C1N)C(=NN2C(C)C)C3NC4=C(C=3)C=C(C=C4)O |
| -0.58 | 0.01 | 1.61 | Flavopiridol HCl | CDK | SelleckChem | S2679 | C1(=C(C(=C2C(=C1O)C(C=C(O2)C3=C(C=CC=C3)Cl)=O)[C@@]4([C@@H](CN(CC4)C)O)[H])O).Cl |
| -0.59 | -0.57 | -0.59 | LYN-1604-HCl, cat. No. HY-101923A | ULK1 agonist | MedChemExpress | HY-101923A | ClC1=CC=C(C(CN2CCN(C(CN(CC(C)C)CC(C)C)=O)CC2)OCC3=CC=C(C=CC=C4)C4=C3)C(Cl)=C1.[H]Cl |
| -0.62 | -0.34 | 2.92 | SNS-032 (BMS-387032) | CDK | SelleckChem | S1145 | C1(=CN=C(O1)CSC2=CN=C(S2)NC(=O)C3CCNCC3)C(C)(C)C |
| -0.62 | -0.18 | 0.89 | Honokiol | MEK,Akt | SelleckChem | S2310 | C1(=C(C=C(C=C1)C2=CC(=CC=C2O)CC=C)CC=C)O |
| -0.62 | -0.57 | 0.94 | XL388 | mTOR | SelleckChem | S7035 | C1(=NC=C(C=C1)C2=CC=C3C(=C2)CN(CCO3)C(=O)C4=CC=C(C(=C4C)F)S(C)(=O)=O)N |
| -0.63 | -0.8 | 2.81 | NVP-BHG712 | Raf,Src,Bcr-Abl,VEGFR,Ephrin receptor | SelleckChem | S2202 | C12=C(N=C(N=C1NC3=CC(=CC=C3C)C(NC4=CC=CC(=C4)C(F)(F)F)=O)C5=CC=CN=C5)N(N=C2)C |
| -0.64 | -0.45 | 0.89 | ZM 336372 | Raf | SelleckChem | S2720 | C1(=CC=CC(=C1)C(NC2=CC(=C(C=C2)C)NC(C3=CC=C(C=C3)O)=O)=O)N(C)C |
| -0.66 | -0.19 | 0.81 | AG-490 (Tyrphostin B42) | JAK,EGFR | SelleckChem | S1143 | C1(=C(C=CC(=C1)/C=C(/C(NCC2=CC=CC=C2)=O)C#N)O)O |
| -0.67 | -0.74 | 2.03 | PD318088 | MEK | SelleckChem | S1568 | C1=C(C(=C(C(=C1C(NOCC(CO)O)=O)NC2=CC=C(C=C2F)I)F)F)Br |
| -0.67 | -0.7 | 0.85 | SKI II | S1P Receptor | SelleckChem | S7176 | C1(C2=CC=C(C=C2)Cl)N=C(NC3=CC=C(C=C3)O)SC=1 |
| -0.68 | -0.56 | 0.68 | Y-27632 2HCl | Autophagy,ROCK | SelleckChem | S1049 | [C@@H]1(CC[C@@H](CC1)[C@@H](C)N)C(=O)NC2(C=CN=CC=2).Cl.Cl |
| -0.68 | -0.65 | 1.46 | TAK-632 | Raf | SelleckChem | S7291 | C12=C(C(=C(C=C1)OC3=CC(=C(C=C3)F)NC(CC4=CC=CC(=C4)C(F)(F)F)=O)C#N)SC(=N2)NC(C5CC5)=O |
| -0.69 | -0.28 | 4.67 | OSI-420 | EGFR | SelleckChem | S2205 | C1(=C(C=CC=C1NC2=NC=NC3=C2C=C(C(=C3)OCCOC)OCCO)C#C).Cl |
| -0.69 | -0.27 | 1.23 | GW5074 | Raf | SelleckChem | S2872 | C1=C2C(=CC=C1I)NC(C/2=C/C3=CC(=C(C(=C3)Br)O)Br)=O |
| -0.72 | -0.7 | 1.16 | Nilotinib (AMN-107) | Bcr-Abl | SelleckChem | S1033 | C1(=CC=C(C(=C1)NC2=NC=CC(=N2)C3=CN=CC=C3)C)C(=O)NC4=CC(=CC(=C4)N5C=C(N=C5)C)C(F)(F)F |
| -0.72 | -0.43 | 0.93 | GSK461364 | PLK | SelleckChem | S2193 | C1N(CCN(C1)CC2=CC3=C(C=C2)N=CN3C4=CC(=C(S4)C(=O)N)O[C@H](C)C5=CC=CC=C5C(F)(F)F)C |
| -0.73 | -0.14 | 0.04 | Rigosertib (ON-01910) | PLK | SelleckChem | S1362 | C1(=C(C=C(C=C1OC)OC)OC)/C=C/S(CC2=CC=C(C(=C2)NCC([O-])=O)OC)(=O)=O.[Na+] |
| -0.73 | -0.41 | 2.42 | H 89 2HCl | PKA | SelleckChem | S1582 | C1(=CC=C(C2=C1C=NC=C2)S(NCCNC/C=C/C3=CC=C(C=C3)Br)(=O)=O).Cl.Cl |
| -0.73 | -0.05 | 2.78 | PP121 | DNA-PK,PDGFR,mTOR | SelleckChem | S2622 | C1(=CN=C2C(=C1)C=CN2)C3=NN(C4=C3C(=NC=N4)N)C5CCCC5 |
| -0.73 | -0.88 | 2.29 | TAK-285 | EGFR,HER2 | SelleckChem | S2784 | C12=C(C(=NC=N1)NC3=CC=C(C(=C3)Cl)OC4=CC=CC(=C4)C(F)(F)F)N(C=C2)CCNC(=O)CC(O)(C)C |
| -0.74 | -0.74 | 1.38 | 10058-F4 | c-Myc | SelleckChem | S7153 | C1=C(C=CC(=C1)/C=C2\C(NC(S2)=S)=O)CC |
| -0.74 | -0.77 | 1.45 | CEP-32496 | CSF-1R,Raf | SelleckChem | S8015 | N(C(=O)NC1=NOC(=C1)C(C(F)(F)F)(C)C)C2C=C(C=CC=2)OC3N=CN=C4C=C(C(=CC=34)OC)OC |
| -0.75 | -0.73 | 0.69 | Selumetinib (AZD6244) | MEK | SelleckChem | S1008 | Cn1cnc2c(c(c(cc12)C(NCOCO)=O)Nc1ccc(cc1[Cl])[Br])F |
| -0.75 | -0.6 | 1.2 | Fasudil (HA-1077) HCl | ROCK,Autophagy | SelleckChem | S1573 | C1(=C2C(=C(C=C1)S(N3CCNCCC3)(=O)=O)C=CN=C2).Cl |
| -0.75 | -0.73 | 0.95 | RAF265 (CHIR-265) | VEGFR,Raf | SelleckChem | S2161 | C12=C(C=CC(=C1)OC3=CC=NC(=C3)C4NC(=CN=4)C(F)(F)F)N(C(=N2)NC5=CC=C(C=C5)C(F)(F)F)C |
| -0.75 | -0.58 | 0.89 | PIK-293 | PI3K | SelleckChem | S2207 | C1(N(C(C2=C(N=1)C=CC=C2C)=O)C3=C(C=CC=C3)C)CN4N=CC5=C4N=CN=C5N |
| -0.75 | -0.4 | 1.17 | VX-702 | p38 MAPK | SelleckChem | S6005 | C1=CC(=C(C(=C1)F)N(C2=NC(=C(C=C2)C(N)=O)C3=C(C=C(C=C3)F)F)C(=O)N)F |
| -0.75 | -0.44 | 2.46 | TCS 359 | FLT3 | SelleckChem | S8023 | C12=C(C(=C(S1)NC(C3C=C(C(=CC=3)OC)OC)=O)C(=O)N)CCCC2 |
| -0.76 | -1.55 | 0.96 | Piceatannol | Syk | SelleckChem | S3026 | C1(C(=CC(=CC=1)/C=C/C2C=C(C=C(C=2)O)O)O)O |
| -0.77 | -0.54 | 2.91 | Vatalanib (PTK787) 2HCl | c-Kit,VEGFR | SelleckChem | S1101 | C1(=CC=C2C(=C1)C(=NN=C2CC3=CC=NC=C3)NC4=CC=C(C=C4)Cl).Cl.Cl |
| -0.78 | -0.26 | 0.62 | GSK1904529A | IGF-1R | SelleckChem | S1093 | C1=C(C(=C(C=C1)F)NC(C2=C(C=CC(=C2)C3=C(N4C(=N3)C=CC=C4)C5=CC=NC(=N5)NC6=C(C=C(C(=C6)CC)N7CCC(CC7)N8CCN(CC8)S(C)(=O)=O)OC)OC)=O)F |
| -0.78 | -0.49 | 0.71 | OSU-03012 (AR-12) | PDK-1 | SelleckChem | S1106 | C12=C(C=CC3=C1C=CC(=C3)C4=CC(=NN4C5=CC=C(C=C5)NC(CN)=O)C(F)(F)F)C=CC=C2 |
| -0.78 | -0.63 | 0.78 | Tyrphostin AG 1296 | FGFR,c-Kit,PDGFR | SelleckChem | S8024 | N1C(=CN=C2C=C(C(=CC=12)OC)OC)C3C=CC=CC=3 |
| -0.8 | -0.38 | 0.83 | PD184352 (CI-1040) | MEK | SelleckChem | S1020 | C1=C(C(=C(C(=C1)C(NOCC2CC2)=O)NC3=CC=C(C=C3Cl)I)F)F |
| -0.8 | -0.68 | 2.02 | BIX 02189 | MEK | SelleckChem | S1531 | C1=C(C=C2C(=C1)/C(C(N2)=O)=C(/NC3=CC(=CC=C3)CN(C)C)C4=CC=CC=C4)C(N(C)C)=O |
| -0.8 | -0.54 | 1.77 | CHIR-99021 (CT99021) HCl | GSK-3 | SelleckChem | S2924 | C1(=NC=C(C(=N1)C2=CC=C(C=C2Cl)Cl)C3NC=C(N=3)C)NCCNC4(=NC=C(C=C4)C#N).Cl |
| -0.81 | -0.43 | 1.18 | Neratinib (HKI-272) | HER2,EGFR | SelleckChem | S2150 | C1(=C(C=C2C(=C1)N=CC(=C2NC3=CC=C(C(=C3)Cl)OCC4=NC=CC=C4)C#N)NC(/C=C/CN(C)C)=O)OCC |
| -0.82 | -0.31 | 1.37 | JNJ-38877605 | c-Met | SelleckChem | S1114 | N1=CC=CC2=C1C=CC(=C2)C(C3N4C(=NN=3)C=CC(=N4)C5C=NN(C=5)C)(F)F |
| -0.82 | -0.42 | 0.73 | Triciribine | Akt | SelleckChem | S1117 | [C@H]1([C@@H]([C@H](O[C@@H]1N2C3=C4C(=C2)C(=NN(C4=NC=N3)C)N)CO)O)O |
| -0.82 | -0.41 | 1.21 | AS-604850 | PI3K | SelleckChem | S2681 | C12=C(C=CC(=C1)/C=C3\SC(NC3=O)=O)OC(O2)(F)F |
| -0.83 | -0.55 | 1.38 | AZD8931 (Sapitinib) | HER2,EGFR | SelleckChem | S2192 | C1(=C(C=C2C(=C1)C(=NC=N2)NC3=C(F)C(=CC=C3)Cl)OC)OC4CCN(CC4)CC(NC)=O |
| -0.83 | -0.33 | 0.91 | CAY10505 | PI3K | SelleckChem | S2682 | C1(=CC=C(C=C1)C2OC(=CC=2)/C=C3\C(NC(S3)=O)=O)F |
| -0.85 | -0.02 | 0.81 | SL-327 | MEK | SelleckChem | S1066 | C1=CC=CC(=C1C(F)(F)F)/C(=C(/SC2=CC=C(C=C2)N)N)C#N |
| -0.85 | -0.47 | 2.86 | Torin 2 | ATM/ATR,mTOR | SelleckChem | S2817 | N1(C(C=CC2C=NC3C(C1=2)=CC(=CC=3)C4C=NC(=CC=4)N)=O)C5C=C(C=CC=5)C(F)(F)F |
| -0.85 | -0.43 | 0.93 | SSR128129E | FGFR | SelleckChem | S7167 | C1=CC2N(C(=C(C=2OC)C)C(=O)C3=CC=C(C(=C3)C(O[Na])=O)N)C=C1 |
| -0.86 | -0.59 | 1.77 | Gefitinib (ZD1839) | EGFR | SelleckChem | S1025 | C1(=C(C=C2C(=C1)N=CN=C2NC3=CC=C(C(=C3)Cl)F)OCCCN4CCOCC4)OC |
| -0.86 | -0.42 | 0.82 | GSK2636771 | PI3K | SelleckChem | S8002 | C1=C(C=C2C(=C1C(=O)O)N=C(N2CC3=CC=CC(=C3C)C(F)(F)F)C)N4CCOCC4 |
| -0.86 | -0.74 | 0.98 | SMI-4a | Pim | SelleckChem | S8005 | S1C(NC(C/1=C\C2C=C(C=CC=2)C(F)(F)F)=O)=O |
| -0.87 | -0.46 | 0.61 | GSK690693 | Akt | SelleckChem | S1113 | C12=C(C(=CN=C1C#CC(C)(O)C)OC[C@@H]3CNCCC3)N(C(=N2)C4C(=NON=4)N)CC |
| -0.87 | -0.93 | 0.86 | Quizartinib (AC220) | FLT3 | SelleckChem | S1526 | C1=C(C=CC(=C1)C2N=C3N(C=2)C4C(S3)=CC(=CC=4)OCCN5CCOCC5)NC(NC6C=C(ON=6)C(C)(C)C)=O |
| -0.87 | -0.15 | 4.02 | KRN 633 | PDGFR,VEGFR | SelleckChem | S1557 | C1(=C(C=C2C(=C1)C(=NC=N2)OC3=CC=C(C(=C3)Cl)NC(=O)NCCC)OC)OC |
| -0.87 | -1.24 | 1.01 | GNF-2 | Bcr-Abl | SelleckChem | S2899 | C1=CC(=CC(=C1)C(=O)N)C2=NC=NC(=C2)NC3=CC=C(C=C3)OC(F)(F)F |
| -0.88 | -0.49 | 1.04 | A-769662 | AMPK | SelleckChem | S2697 | C1(=CC=C(C=C1)C2=CSC3=C2C(=C(C(N3)=O)C#N)O)C4=CC=CC=C4O |
| -0.89 | -0.55 | 1.98 | Go 6983 | PKC | SelleckChem | S2911 | C1(=CC=C2C(=C1)C(=CN2CCCN(C)C)C3C(NC(C=3C4C5=C(C=CC=C5)NC=4)=O)=O)OC |
| -0.89 | -0.43 | 1.35 | Icotinib | EGFR | SelleckChem | S2922 | C1=C(C=CC=C1NC2=NC=NC3=C2C=C4C(=C3)OCCOCCOCCO4)C#C |
| -0.9 | -0.37 | 6.47 | ZM 447439 | Aurora Kinase | SelleckChem | S1103 | O1CCN(CC1)CCCOC2=C(C=C3C(=C2)N=CN=C3NC4=CC=C(C=C4)NC(=O)C5=CC=CC=C5)OC |
| -0.9 | -0.37 | 0.27 | ETP-46464 | mTOR,ATM/ATR | SelleckChem | S8050 | C1=CC(=CC2=C1N=CC3=C2N(C(OC3)=O)C4=CC=C(C=C4)C(C#N)(C)C)C5=CN=C6C(=C5)C=CC=C6 |
| -0.91 | -0.55 | 0.75 | Acadesine | AMPK | SelleckChem | S1802 | [C@@H]1([C@H]([C@@H](O[C@H]1CO)N2C=NC(=C2N)C(=O)N)O)O |
| -0.91 | -0.47 | 0.98 | BIO | GSK-3 | SelleckChem | S7198 | C1=CC=C2C(=C1)C(\C(N2)=C3\C(NC4=C3C=CC(=C4)Br)=O)=N\O |
| -0.91 | -0.5 | 0.88 | EHop-016 | Rac | SelleckChem | S7319 | C1(N=C(C=CN=1)NC2C=CC3N(C4C=CC=CC(C(C=2)=3)=4)CC)NCCCN5CCOCC5 |
| -0.92 | -0.36 | 1.41 | CP-724714 | EGFR,HER2 | SelleckChem | S1167 | C1(=CC=C2C(=C1)C(=NC=N2)NC3=CC=C(C(=C3)C)OC4=CC=C(N=C4)C)/C=C/CNC(COC)=O |
| -0.92 | -0.72 | 7.63 | Pazopanib | PDGFR,c-Kit,VEGFR | SelleckChem | S3012 | C1(C(=CC=C(C=1)NC2N=C(C=CN=2)N(C)C3C=CC4=C(N(N=C(C=3)4)C)C)C)S(=O)(=O)N |
| -0.93 | -0.05 | 1.04 | Enzastaurin (LY317615) | PKC | SelleckChem | S1055 | C1=CC=C2C(=C1)C(=CN2C3CCN(CC3)CC4=NC=CC=C4)C5C(NC(C=5C6C7C(N(C=6)C)=CC=CC=7)=O)=O |
| -0.93 | -0.94 | 1.06 | ZM 323881 HCl | VEGFR | SelleckChem | S2896 | C1(C(=CC(=C(C=1)NC2N=CN=C3C=C(C=CC=23)OCC4C=CC=CC=4)F)C)O.Cl |
| -0.93 | -1.01 | 0.66 | IPI-145 (INK1197) | PI3K | SelleckChem | S7028 | C1=CC=C2C(=C1Cl)C(N(C(=C2)[C@H](C)NC3=NC=NC4=C3N=CN4)C5=CC=CC=C5)=O |
| -0.94 | -0.4 | 1.65 | BIX 02188 | MEK | SelleckChem | S1530 | C1=C(C=C2C(=C1)/C(C(N2)=O)=C(/NC3=CC(=CC=C3)CN(C)C)C4=CC=CC=C4)C(NC)=O |
| -0.95 | -0.31 | 5.29 | Lenvatinib (E7080) | VEGFR | SelleckChem | S1164 | C1(=C(C=C2C(=C1)C(=CC=N2)OC3=CC(=C(C=C3)NC(=O)NC4CC4)Cl)OC)C(N)=O |
| -0.95 | -0.53 | 0.99 | CZC24832 | PI3K | SelleckChem | S7018 | C1(=CN=CC(=C1)C2C=C(C3=NC(=NN(C=2)3)N)F)S(=O)(=O)NC(C)(C)C |
| -0.96 | -0.48 | 1.95 | Motesanib Diphosphate (AMG-706) | VEGFR,PDGFR,c-Kit | SelleckChem | S1032 | C1(=CN=CC=C1CNC2=C(C=CC=N2)C(=O)NC3=CC=C4C(=C3)NCC4(C)C).P(O)(=O)(O)O.P(O)(=O)(O)O |
| -0.96 | -0.23 | 0.5 | LY294002 | Autophagy,PI3K | SelleckChem | S1105 | C1=CC(=C2C(=C1)C(C=C(O2)N3CCOCC3)=O)C4=CC=CC=C4 |
| -0.96 | -0.51 | 0.72 | SGX-523 | c-Met | SelleckChem | S1112 | C1=C2C(=CC(=C1)SC3N4C(=NN=3)C=CC(=N4)C5C=NN(C=5)C)C=CC=N2 |
| -0.96 | -0.45 | 1.54 | Brivanib Alaninate (BMS-582664) | VEGFR,FGFR | SelleckChem | S1138 | C12=C(C(=C(C=C1)OC3=NC=NN4C3=C(C(=C4)OC[C@@H](C)OC([C@@H](N)C)=O)C)F)C=C(N2)C |
| -0.96 | -0.51 | 0.77 | NSC 23766 | Rac | SelleckChem | S8031 | N1(C(=CC(=C2C=C(C=CC=12)NC3=NC(=NC(=C3)C)NC(CCCN(CC)CC)C)N)C).Cl.Cl.Cl |
| -0.97 | -0.56 | 1.55 | PD0325901 | MEK | SelleckChem | S1036 | C1=CC(=C(C(=C1C(=O)NOC[C@H](O)CO)NC2=CC=C(C=C2F)I)F)F |
| -0.97 | -0.62 | 0.85 | NU7026 | DNA-PK | SelleckChem | S2893 | C1(OC2=C(C=CC3=C2C=CC=C3)C(C=1)=O)N4CCOCC4 |
| -0.98 | -0.2 | 1.42 | Cediranib (AZD2171) | VEGFR | SelleckChem | S1017 | C1(=C(C=C2C(=C1)C(=NC=N2)OC3=C(C4=C(C=C3)NC(=C4)C)F)OCCCN5CCCC5)OC |
| -0.98 | -0.45 | 0.72 | XL147 | PI3K | SelleckChem | S1118 | C1=CC=C2C(=C1)N=C(C(=N2)NS(=O)(=O)C3=CC=C(C=C3)C)NC4=CC5C(C=C4)=NSN=5 |
| -0.98 | -0.52 | 5.04 | GSK429286A | ROCK | SelleckChem | S1474 | C1(CC(C(=C(N1)C)C(NC2=C(C=C3C(=C2)C=NN3)F)=O)C4=CC=C(C=C4)C(F)(F)F)=O |
| -0.98 | -0.66 | 1.09 | AS-252424 | PI3K | SelleckChem | S2671 | C1(=CC(=C(C=C1)C2OC(=CC=2)/C=C3\SC(NC3=O)=O)O)F |
| -0.98 | -0.35 | 0.69 | INCB28060 | c-Met | SelleckChem | S2788 | C(C1C(=CC(=CC=1)C2C=NC3N(N=2)C(=CN=3)CC4C=C5C=CC=NC(=CC=4)5)F)(=O)NC |
| -0.98 | -0.37 | 1.16 | TG100713 | PI3K | SelleckChem | S2870 | C1(C=C(C=CC=1)C2=NC3=C(N=C(N=C(N=C2)3)N)N)O |
| -0.98 | -0.9 | 1.62 | SC-514 | IκB/IKK | SelleckChem | S4907 | C1(=CC(=C(S1)C(=O)N)N)C2=CSC=C2 |
| -0.98 | -0.62 | 0.74 | P276-00 | CDK | SelleckChem | S8058 | C1(=CC(=C(C=C1)C2=CC(C3=C(O2)C(=C(C=C3O)O)[C@H]4[C@@H](N(CC4)C)CO)=O)Cl).Cl |
| -0.99 | -0.56 | 1.56 | AZD8055 | mTOR | SelleckChem | S1555 | C1(=C(C=CC(=C1)C2=NC3=C(C=C2)C(=NC(=N3)N4[C@H](COCC4)C)N5[C@H](COCC5)C)OC)CO |
| -0.99 | -0.59 | 0.6 | Ridaforolimus (Deforolimus, MK-8669) | mTOR | SelleckChem | S1022 | [C@H]1(C[C@@H](CC[C@@H]1OP(C)(=O)C)C[C@H]([C@@H]2CC([C@@H](/C=C(/[C@H]([C@@H](OC)C([C@@H](C[C@@H](/C=C/C=C/C=C(/[C@H](C[C@]3(CC[C@H]([C@](C(C(N4[C@](C(O2)=O)(CCCC4)[H])=O)=O)(O3)O)C)[H])OC)C)C)C)=O)O)C)C)=O)C)OC |
| -0.99 | -0.48 | 2.57 | ZM 306416 | VEGFR | SelleckChem | S2897 | C12C(C=C(C(=C1)OC)OC)=NC=NC=2NC3=C(C=C(C=C3)Cl)F |
| -1 | -0.46 | 4.79 | Erlotinib HCl (OSI-744) | Autophagy,EGFR | SelleckChem | S1023 | C1(=CC=CC(=C1)C#C)NC2(=NC=NC3=C2C=C(C(=C3)OCCOC)OCCOC)C.Cl |
| -1 | -0.44 | 0.86 | Rapamycin (Sirolimus) | Autophagy,mTOR | SelleckChem | S1039 | [C@@H]1([C@@H](C[C@@H](CC1)C[C@H]([C@@H]2CC([C@@H](/C=C(\C)[C@H]([C@@H](C([C@@H](C[C@@H](/C=C/C=C/C=C(/[C@H](C[C@@H]3CC[C@H]([C@](C(C(N4CCCC[C@@H](C(O2)=O)4)=O)=O)(O3)O)C)OC)C)C)C)=O)OC)O)C)=O)C)OC)O |
| -1.01 | -0.41 | 0.66 | MK-2206 2HCl | Akt | SelleckChem | S1078 | C1(=CC=CC=C1)C2(=C(N=C3C(=C2)C4N(C=C3)C(NN=4)=O)C5=CC=C(C=C5)C6(CCC6)N).Cl.Cl |
| -1.01 | 0.11 | 1.09 | GDC-0980 (RG7422) | mTOR,PI3K | SelleckChem | S2696 | C1(=CN=C(N=C1)N)C2=NC(=C3C(=N2)C(=C(S3)CN4CCN(CC4)C([C@@H](O)C)=O)C)N5CCOCC5 |
| -1.02 | -0.46 | 0.69 | TG003 | CDK | SelleckChem | S7320 | C12N(/C(=C/C(C)=O)SC=1C=CC(=C2)OC)CC |
| -1.03 | -0.7 | 0.84 | Everolimus (RAD001) | mTOR | SelleckChem | S1120 | C(O)CO[C@H]1[C@@H](C[C@@H](CC1)C[C@@H]([C@H]2OC(=O)[C@@H]3CCCCN3C(=O)C([C@]4([C@@H](CC[C@H](O4)C[C@@H](/C(=C/C=C/C=C/[C@H](C[C@H](C([C@@H]([C@@H](O)/C(=C/[C@H](C(C2)=O)C)C)OC)=O)C)C)C)OC)C)O)=O)C)OC |
| -1.03 | -0.59 | 1.68 | Varlitinib | EGFR | SelleckChem | S2755 | C12=C(C(=NC=N1)NC3=CC(=C(C=C3)OCC4SC=CN=4)Cl)C=C(C=C2)NC5OC[C@H](N=5)C |
| -1.04 | -0.6 | 20.74 | MLN8054 | Aurora Kinase | SelleckChem | S1100 | C1=C(C=C2C(=C1)C3=C(CN=C2C4=C(C=CC=C4F)F)C=NC(=N3)NC5=CC=C(C=C5)C(=O)O)Cl |
| -1.04 | -0.66 | 3.43 | Thiazovivin | ROCK | SelleckChem | S1459 | C1(SC=C(N=1)C(NCC2=CC=CC=C2)=O)NC3=CC=NC=N3 |
| -1.04 | -0.46 | 2.05 | SGI-1776 free base | Pim | SelleckChem | S2198 | C1C(=NN2C(C=1)=NC=C2C3=CC=CC(=C3)OC(F)(F)F)NCC4CCN(CC4)C |
| -1.05 | -0.5 | 0.88 | Amuvatinib (MP-470) | FLT3,c-RET,PDGFR,c-Kit | SelleckChem | S1244 | C1=CC=C2C(=C1)C3=C(O2)C(=NC=N3)N4CCN(CC4)C(=S)NCC5=CC=C6C(=C5)OCO6 |
| -1.06 | -0.5 | 1.98 | GDC-0941 | PI3K | SelleckChem | S1065 | C1(=CC=CC2=C1C=NN2)C3=NC(=C4C(=N3)C=C(S4)CN5CCN(CC5)S(=O)(=O)C)N6CCOCC6 |
| -1.06 | -0.51 | 0.82 | ZCL278 | Rac | SelleckChem | S7293 | C1=C(N=C(N=C1C)NS(C2=CC=C(C=C2)NC(NC(COC3=C(C=C(C=C3)Br)Cl)=O)=S)(=O)=O)C |
| -1.07 | -0.43 | 0.59 | ZSTK474 | PI3K | SelleckChem | S1072 | N1=C(N=C(N=C1N2CCOCC2)N3C4=C(N=C3C(F)F)C=CC=C4)N5CCOCC5 |
| -1.07 | -0.58 | 2.45 | SP600125 | JNK | SelleckChem | S1460 | C1C=CC2=C(C=1)C(C3=C4C2=NNC4=CC=C3)=O |
| -1.07 | -0.28 | 1.52 | CH5132799 | mTOR,PI3K | SelleckChem | S2699 | N1=C(N=CC(=C1)C2=C3C(=NC(=N2)N4CCOCC4)N(CC3)S(C)(=O)=O)N |
| -1.07 | -0.58 | 2.15 | INK 128 (MLN0128) | mTOR | SelleckChem | S2811 | N1=CN=C2C(=C1N)C(=NN2C(C)C)C3=CC4=C(C=C3)OC(=N4)N |
| -1.07 | -0.09 | 1.03 | AG-18 | EGFR | SelleckChem | S8009 | C1(C=C(C#N)C#N)=CC(=C(C=C1)O)O |
| -1.08 | -0.91 | 0.72 | GDC-0349 | mTOR | SelleckChem | S8040 | N(C(=O)NC1C=CC(=CC=1)C2N=C3C(=C(N=2)N4[C@H](COCC4)C)CCN(C3)C5COC5)CC |
| -1.09 | -0.56 | 1.22 | Vandetanib (ZD6474) | VEGFR | SelleckChem | S1046 | C12=C(C(=NC=N1)NC3=C(C=C(C=C3)Br)F)C=C(C(=C2)OCC4CCN(CC4)C)OC |
| -1.09 | -0.68 | 1.1 | SB203580 | p38 MAPK | SelleckChem | S1076 | N1=CC=C(C=C1)C2=C(N=C(N2)C3=CC=C(C=C3)S(C)=O)C4=CC=C(C=C4)F |
| -1.09 | -0.53 | 1.11 | GDC-0068 | Akt | SelleckChem | S2808 | C1CN(CCN1C([C@@H](C2=CC=C(C=C2)Cl)CNC(C)C)=O)C3=C4C(=NC=N3)[C@H](C[C@H]4C)O |
| -1.1 | -0.61 | 0.91 | Pimasertib (AS-703026) | MEK | SelleckChem | S1475 | C1=C(C=CC(=C1F)NC2=CN=CC=C2C(NC[C@@H](CO)O)=O)I |
| -1.1 | -0.33 | 0.64 | PF-04217903 | c-Met | SelleckChem | S1094 | C12=CC=CN=C1C=CC(=C2)CN3C4=C(N=N3)N=CC(=N4)C5=CN(N=C5)CCO |
| -1.1 | -0.2 | 0.87 | PH-797804 | p38 MAPK | SelleckChem | S2726 | C1(=C(C=CC(=C1)C(=O)NC)C)N2C(=CC(=C(C2=O)Br)OCC3=CC=C(C=C3F)F)C |
| -1.1 | -0.38 | 1.26 | AZD2014 | mTOR | SelleckChem | S2783 | C1=C(N=C2C(=C1)C(=NC(=N2)N3[C@H](COCC3)C)N4[C@H](COCC4)C)C5=CC(=CC=C5)C(NC)=O |
| -1.1 | -0.37 | 0.69 | CGK 733 | ATM/ATR | SelleckChem | S7136 | C1=C(C(=CC(=C1)NC(NC(C(Cl)(Cl)Cl)NC(C(C2=CC=CC=C2)C3=CC=CC=C3)=O)=S)[N+](=O)[O-])F |
| -1.1 | -1.18 | 1.18 | Skepinone-L | p38 MAPK | SelleckChem | S7214 | C12C(C(C3C(CC1)=CC=C(C=3)OC[C@@H](CO)O)=O)=CC=C(C=2)NC4C(=CC(=CC=4)F)F |
| -1.1 | -0.41 | 0.99 | AR-A014418 | GSK-3 | SelleckChem | S7435 | C1=C(SC(=N1)NC(NCC2=CC=C(C=C2)OC)=O)[N+](=O)[O-] |
| -1.11 | -0.49 | 0.59 | PD98059 | MEK | SelleckChem | S1177 | C1=CC=C2C(=C1)C(C=C(O2)C3=CC=CC(=C3N)OC)=O |
| -1.12 | -0.59 | 0.77 | BEZ235 (NVP-BEZ235, Dactolisib) | PI3K,ATM/ATR,mTOR | SelleckChem | S1009 | C12=CN=C3C(=C1N(C(N2C)=O)C4=CC=C(C=C4)C(C#N)(C)C)C=C(C=C3)C5=CC6=C(N=C5)C=CC=C6 |
| -1.12 | -0.63 | 2.54 | Brivanib (BMS-540215) | FGFR,VEGFR | SelleckChem | S1084 | C12=C(C(=C(C=C1)OC3=NC=NN4C3=C(C(=C4)OC[C@@H](C)O)C)F)C=C(N2)C |
| -1.12 | -0.48 | 1.94 | TG100-115 | PI3K | SelleckChem | S1352 | C1(=C(N=C2C(=N1)C(=NC(=N2)N)N)C3=CC=CC(=C3)O)C4=CC(=CC=C4)O |
| -1.12 | -0.86 | 5.47 | MGCD-265 | Tie-2,VEGFR,c-Met | SelleckChem | S1361 | C1(=CC=CC=C1)CC(NC(NC2=CC=C(C(=C2)F)OC3=C4C(=NC=C3)C=C(S4)C5N=CN(C=5)C)=S)=O |
| -1.12 | -0.54 | 1 | LY2228820 | p38 MAPK | SelleckChem | S1494 | CC(C1=NC(=C(N1)C2=NC3=C(C=C2)N=C(N3CC(C)(C)C)N)C4=CC=C(C=C4)F)(C)C.CS(=O)(=O)O.CS(=O)(=O)O |
| -1.13 | -0.45 | 3.59 | AT7867 | S6 Kinase,Akt | SelleckChem | S1558 | C1(=CC=C(C=C1)C2(C3=CC=C(C=C3)C4=CNN=C4)CCNCC2)Cl |
| -1.13 | -0.53 | 0.75 | AZD8330 | MEK | SelleckChem | S2134 | C1=C(C=CC(=C1F)NC2N(C(C(=CC=2C(NOCCO)=O)C)=O)C)I |
| -1.13 | -0.67 | 1.13 | TAK-715 | p38 MAPK | SelleckChem | S2928 | C(C1C=CC=CC=1)(=O)NC2C=C(C=CN=2)C3=C(N=C(S3)CC)C4C=C(C=CC=4)C |
| -1.14 | -0.48 | 0.74 | KU-55933 (ATM Kinase Inhibitor) | ATM/ATR | SelleckChem | S1092 | C1=CC=C2C(=C1)SC3=C(S2)C=CC=C3C4=CC(C=C(O4)N5CCOCC5)=O |
| -1.14 | -0.19 | 0.9 | PF-04691502 | Akt,mTOR,PI3K | SelleckChem | S2743 | N1=C(N=C2C(=C1C)C=C(C(N2[C@H]3CC[C@@H](CC3)OCCO)=O)C4=CC=C(N=C4)OC)N |
| -1.14 | -0.71 | 0.93 | CUDC-907 | PI3K,HDAC | SelleckChem | S2759 | C1=NC(=NC=C1C(NO)=O)N(C)CC2=CC3=C(S2)C(=NC(=N3)C4=CC=C(N=C4)OC)N5CCOCC5 |
| -1.14 | -0.66 | 1.85 | Tofacitinib (CP-690550) Citrate | JAK | SelleckChem | S5001 | N1(=CN=C2C(=C1N(C)[C@H]3CN(CC[C@@H]3C)C(=O)CC#N)C=CN2).C(C(C(O)=O)(CC(=O)O)O)C(O)=O |
| -1.15 | -0.71 | 1.53 | WYE-354 | mTOR | SelleckChem | S1266 | C1=C(C=CC(=C1)C2=NC3=C(C(=N2)N4CCOCC4)C=NN3C5CCN(CC5)C(=O)OC)NC(OC)=O |
| -1.16 | 0.01 | 0.85 | WYE-125132 (WYE-132) | mTOR | SelleckChem | S2661 | C1(=NC(=NC2=C1C=NN2C3CCC4(CC3)OCCO4)C5=CC=C(C=C5)NC(=O)NC)N6CC7CCC(O7)C6 |
| -1.16 | -0.68 | 1.02 | NVP-BVU972 | c-Met | SelleckChem | S2761 | C1C(=NN2C(C=1)=NC=C2CC3=CC=C4C(=C3)C=CC=N4)C5C=NN(C=5)C |
| -1.16 | -0.35 | 0.64 | Alectinib (CH5424802) | ALK | SelleckChem | S2762 | C1(=CC=C2C(=C1)NC3=C2C(C4=C(C3(C)C)C=C(C(=C4)CC)N5CCC(CC5)N6CCOCC6)=O)C#N |
| -1.17 | -0.55 | 1.97 | Tandutinib (MLN518) | FLT3 | SelleckChem | S1043 | C1=C(C(=CC2=C1N=CN=C2N3CCN(CC3)C(NC4=CC=C(C=C4)OC(C)C)=O)OC)OCCCN5CCCCC5 |
| -1.17 | -0.77 | 25.69 | MK-5108 (VX-689) | Aurora Kinase | SelleckChem | S2770 | N1=C(C=CC=C1C[C@]2(CC[C@H](CC2)OC3=C(C(=CC=C3)Cl)F)C(=O)O)NC4SC=CN=4 |
| -1.18 | -0.39 | 3.53 | Canertinib (CI-1033) | EGFR,HER2 | SelleckChem | S1019 | C1(=C(C=C2C(=C1)C(=NC=N2)NC3=CC(=C(C=C3)F)Cl)OCCCN4CCOCC4)NC(C=C)=O |
| -1.18 | -0.65 | 0.88 | PI-103 | PI3K,Autophagy,DNA-PK,mTOR | SelleckChem | S1038 | C12=C(N=C(N=C1N3CCOCC3)C4=CC=CC(=C4)O)C5=C(O2)N=CC=C5 |
| -1.18 | -0.73 | 0.77 | Temsirolimus (CCI-779, NSC 683864) | mTOR | SelleckChem | S1044 | [C@H]1(C[C@@H](CC[C@@H]1OC(C(CO)(CO)C)=O)C[C@H]([C@@H]2CC([C@@H](/C=C(/[C@H]([C@@H](OC)C([C@@H](C[C@@H](/C=C/C=C/C=C(/[C@H](C[C@]3(CC[C@H]([C@](C(C(N4[C@](C(O2)=O)(CCCC4)[H])=O)=O)(O3)O)C)[H])OC)C)C)C)=O)O)C)C)=O)C)OC |
| -1.18 | -0.52 | 1.1 | BI 2536 | PLK | SelleckChem | S1109 | C1(=CC=C(C(=C1)OC)NC2=NC=C3C(=N2)N([C@@H](C(N3C)=O)CC)C4CCCC4)C(NC5CCN(CC5)C)=O |
| -1.18 | -0.5 | 1.08 | PD168393 | EGFR | SelleckChem | S7039 | C1=C(C=C2C(=C1)N=CN=C2NC3=CC=CC(=C3)Br)NC(C=C)=O |
| -1.19 | -0.65 | 7.78 | Pazopanib HCl (GW786034 HCl) | VEGFR,PDGFR,c-Kit | SelleckChem | S1035 | C1(=C(C(=CC(=C1)NC2=NC=CC(=N2)N(C3C=CC4C(C=3)=NN(C=4C)C)C)S(=O)(N)=O)C).Cl |
| -1.19 | -0.34 | 0.93 | AC480 (BMS-599626) | HER2,EGFR | SelleckChem | S1056 | C1(CN[C@@H](CO1)COC(NC2=CN3C(=C2C)C(=NC=N3)NC4=CC5=C(C=C4)N(N=C5)CC6=CC(=CC=C6)F)=O).Cl |
| -1.19 | -0.36 | 1.07 | SB202190 (FHPI) | p38 MAPK | SelleckChem | S1077 | C1=C(C=CC(=C1)C2N=C(NC=2C3=CC=NC=C3)C4=CC=C(C=C4)O)F |
| -1.19 | -0.67 | 1.16 | WAY-600 | mTOR | SelleckChem | S2689 | C12=C(C=CC(=C1)C3=NC4=C(C(=N3)N5CCOCC5)C=NN4C6CCN(CC6)CC7=CC=CN=C7)NC=C2 |
| -1.2 | -0.31 | 0.99 | BAY 11-7082 | IκB/IKK,E2 conjugating | SelleckChem | S2913 | C1(S(/C=C/C#N)(=O)=O)C=CC(=CC=1)C |
| -1.22 | -0.54 | 0.97 | Zoledronic Acid | Rac | SelleckChem | S1314 | C1=NC=CN1CC(O)(P(=O)(O)O)P(=O)(O)O |
| -1.22 | -0.72 | 3.2 | PF-4708671 | S6 Kinase | SelleckChem | S2163 | N1C(=NC2=C1C=CC(=C2)C(F)(F)F)CN3CCN(C4=NC=NC=C4CC)CC3 |
| -1.22 | -0.5 | 1.07 | AZ20 | ATM/ATR | SelleckChem | S7050 | C1=C(N=C(N=C1N2[C@@H](COCC2)C)C3=CC=CC4=C3C=CN4)C5(S(C)(=O)=O)CC5 |
| -1.24 | -0.4 | 1.55 | Tofacitinib (CP-690550,Tasocitinib) | JAK | SelleckChem | S2789 | N1=CN=C2C(=C1N(C)[C@H]3CN(CC[C@@H]3C)C(=O)CC#N)C=CN2 |
| -1.25 | -0.21 | 1.62 | AG-1478 (Tyrphostin AG-1478) | EGFR | SelleckChem | S2728 | C1=C2C(=CC(=C1OC)OC)C(=NC=N2)NC3=CC(=CC=C3)Cl |
| -1.25 | -0.56 | 0.77 | 3-Methyladenine | Autophagy,PI3K | SelleckChem | S2767 | N1=CN=C2N(C=NC(=C12)N)C |
| -1.26 | -0.51 | 0.88 | A66 | PI3K | SelleckChem | S2636 | C1SC(=NC=1C2SC(=NC=2C)NC(N3[C@@H](CCC3)C(=O)N)=O)C(C)(C)C |
| -1.26 | -0.48 | 1.81 | BGT226 (NVP-BGT226) | PI3K,mTOR | SelleckChem | S2749 | C(/C=C\C(=O)O)(=O)O.N1=CC2=C(C3=CC(=CC=C13)C4C=NC(=CC=4)OC)N(C(N2C)=O)C5C=C(C(=CC=5)N6CCNCC6)C(F)(F)F |
| -1.26 | -0.44 | 0.93 | TIC10 | Akt | SelleckChem | S7127 | C1=CC=CC(=C1)CN2CCC3=C(C2)C(N4C(N3CC5=C(C=CC=C5)C)=NCC4)=O |
| -1.27 | -0.44 | 0.82 | Mubritinib (TAK 165) | HER2 | SelleckChem | S2216 | C1=C(C=CC(=C1)/C=C/C2OC=C(N=2)COC3=CC=C(C=C3)CCCCN4N=NC=C4)C(F)(F)F |
| -1.27 | -0.78 | 1.16 | Torin 1 | Autophagy,mTOR | SelleckChem | S2827 | N1(C(C=CC2C=NC3C(C1=2)=CC(=CC=3)C4C=NC5=CC=CC=C(C=4)5)=O)C6C=C(C(=CC=6)N7CCN(CC7)C(CC)=O)C(F)(F)F |
| -1.29 | -0.65 | 0.71 | KU-60019 | ATM/ATR | SelleckChem | S1570 | C1=C(C=CC2=C1CC3=C(S2)C(=CC=C3)C4=CC(C=C(O4)N5CCOCC5)=O)NC(CN6C[C@@H](O[C@@H](C6)C)C)=O |
| -1.29 | -0.46 | -0.9 | IPA-3 | PAK | SelleckChem | S7093 | S(SC1C(=CC=C2C=CC=CC=12)O)C3C(=CC=C4C=CC=CC=34)O |
| -1.3 | -0.39 | 1.18 | PD173074 | VEGFR,FGFR | SelleckChem | S1264 | N1=C(N=C2C(=C1)C=C(C(=N2)NC(NC(C)(C)C)=O)C3=CC(=CC(=C3)OC)OC)NCCCCN(CC)CC |
| -1.3 | -0.6 | 0.9 | VX-745 | p38 MAPK | SelleckChem | S1458 | C1=C(C=CC(=C1F)SC2C=CC3N(N=2)C=NC(C=3C4=C(C=CC=C4Cl)Cl)=O)F |
| -1.3 | -0.7 | 1.2 | Tie2 kinase inhibitor | Tie-2 | SelleckChem | S1577 | C12=C(C=CC(=C1)C3=C(N=C(N3)C4=CC=C(C=C4)S(C)=O)C5=CC=NC=C5)C=C(C=C2)OC |
| -1.3 | -0.68 | 19.95 | MK-8745 | Aurora Kinase | SelleckChem | S7065 | C1(=CC=CC(=C1F)C(N2CCN(CC2)CC3=CC=CC(=N3)NC4=NC=CS4)=O)Cl |
| -1.3 | -0.87 | 1.98 | RKI-1447 | ROCK | SelleckChem | S7195 | C1=NC=CC(=C1)C2N=C(SC=2)NC(NCC3=CC=CC(=C3)O)=O |
| -1.31 | -0.85 | 2.81 | CUDC-101 | HDAC,HER2,EGFR | SelleckChem | S1194 | C12=C(N=CN=C1NC3=CC(=CC=C3)C#C)C=C(C(=C2)OCCCCCCC(NO)=O)OC |
| -1.31 | -0.43 | 1.43 | ZM 39923 HCl | JAK | SelleckChem | S8004 | C(CCN(C(C)C)CC1C=CC=CC=1)(=O)C2(C=C3C=CC=CC(=CC=2)3).Cl |
| -1.31 | -0.56 | 0.79 | PF-05212384 (PKI-587) | mTOR,PI3K | SelleckChem | S2628 | N1=C(N=C(N=C1N2CCOCC2)C3=CC=C(C=C3)NC(NC4=CC=C(C=C4)C(N5CCC(CC5)N(C)C)=O)=O)N6CCOCC6 |
| -1.31 | 0.3 | 0.81 | CHIR-98014 | GSK-3 | SelleckChem | S2745 | C1=C(C=C(C(=C1)C2=NC(=NC=C2N3C=CN=C3)NCCNC4=CC=C(C(=N4)N)[N+](=O)[O-])Cl)Cl |
| -1.31 | -0.47 | 1.09 | BYL719 | PI3K | SelleckChem | S2814 | N1([C@@H](CCC1)C(=O)N)C(=O)NC2SC(=C(N=2)C)C3=CC(=NC=C3)C(C(F)(F)F)(C)C |
| -1.32 | -0.3 | 1.65 | Imatinib Mesylate (STI571) | c-Kit,Bcr-Abl,PDGFR | SelleckChem | S1026 | C1(=CC=C(C(=C1)NC2=NC=CC(=N2)C3=CC=CN=C3)C)NC(=O)C4(=CC=C(C=C4)CN5CCN(CC5)C).OS(C)(=O)=O |
| -1.32 | -0.63 | 1.05 | AZD6482 | PI3K | SelleckChem | S1462 | C(C1C(=CC=CC=1)N[C@H](C)C2=CC(=CN3C(C=C(N=C32)N4CCOCC4)=O)C)(=O)O |
| -1.32 | -0.22 | 1.29 | LY2603618 | Chk | SelleckChem | S2626 | C1=C(C(=CC(=C1OC[C@@H]2CNCCO2)NC(NC3=CN=C(C=N3)C)=O)Br)C |
| -1.34 | -0.28 | 0.69 | CAL-101 (Idelalisib, GS-1101) | PI3K | SelleckChem | S2226 | C1=CC=C2C(=C1F)C(N(C(=N2)[C@@H](NC3=C4C(=NC=N3)NC=N4)CC)C5=CC=CC=C5)=O |
| -1.36 | -1.21 | 0.42 | Genistein | Topoisomerase,EGFR | SelleckChem | S1342 | C1(=CC2=C(C(=C1)O)C(C(=CO2)C3=CC=C(C=C3)O)=O)O |
| -1.36 | -0.27 | 3.61 | OSI-027 | mTOR | SelleckChem | S2624 | N1C=NN2C(C=1N)=C(N=C2[C@@H]3CC[C@H](CC3)C(=O)O)C4NC5=C(C=4)C=CC=C5OC |
| -1.38 | -0.33 | 1.18 | GSK2126458 (GSK458) | PI3K,mTOR | SelleckChem | S2658 | C1(C(=CC(=CC=1)F)F)S(=O)(=O)NC2C(=NC=C(C=2)C3C=C4C(=CC=NC(=CC=3)4)C5C=NN=CC=5)OC |
| -1.4 | -0.44 | 0.64 | Saracatinib (AZD0530) | Src,Bcr-Abl | SelleckChem | S1006 | C1=C(C=C2C(=C1OC3CCOCC3)C(=NC=N2)NC4=C(C=CC5=C4OCO5)Cl)OCCN6CCN(CC6)C |
| -1.4 | -0.02 | 1.73 | PP2 | Src | SelleckChem | S7008 | N1=CN=C2C(=C1N)C(=NN2C(C)(C)C)C3=CC=C(C=C3)Cl |
| -1.41 | -0.39 | 0.85 | Imatinib (STI571) | PDGFR | SelleckChem | S2475 | N1=C(N=C(C=C1)C2=CC=CN=C2)NC3=CC(=CC=C3C)NC(=O)C4=CC=C(C=C4)CN5CCN(CC5)C |
| -1.41 | -0.49 | 1.36 | SAR131675 | VEGFR | SelleckChem | S2842 | C12C(N(C(=C(C1=O)C(NC)=O)N)CC)=NC(=CC=2)C#C[C@](O)(C)COC |
| -1.41 | -0.46 | 0.9 | MEK162 (ARRY-162, ARRY-438162) | MEK | SelleckChem | S7007 | C12=C(C=C(C(=C1F)NC3=C(C=C(C=C3)Br)F)C(NOCCO)=O)N(C=N2)C |
| -1.42 | -0.45 | 1.21 | PIK-93 | PI3K | SelleckChem | S1489 | C1(=C(C=C(C=C1)C2SC(=NC=2C)NC(C)=O)S(=O)(=O)NCCO)Cl |
| -1.42 | -0.5 | 0.84 | Telatinib | VEGFR,PDGFR,c-Kit | SelleckChem | S2231 | N1=C(C2=C(C(=N1)OCC3=CC(=NC=C3)C(=O)NC)OC=C2)NC4=CC=C(C=C4)Cl |
| -1.42 | -0.85 | 0.79 | Palomid 529 (P529) | mTOR | SelleckChem | S2238 | C1=C(C=CC(=C1)COC2=C(C=C3C(=C2)OC(C4=C3C=CC(=C4)C(C)O)=O)OC)OC |
| -1.43 | 0.52 | 2.06 | Indirubin | GSK-3 | SelleckChem | S2386 | C1=CC=C2C(=C1)NC(C/2=C3\NC4=C(C3=O)C=CC=C4)=O |
| -1.44 | 0.33 | 1.34 | Sotrastaurin | PKC | SelleckChem | S2791 | N1C(C(=C(C1=O)C2N=C(N=C3C=CC=CC=23)N4CCN(CC4)C)C5=CNC6C=CC=CC5=6)=O |
| -1.46 | -0.45 | 0.83 | PHT-427 | PDK-1,Akt | SelleckChem | S1556 | C1=C(C=CC(=C1)S(=O)(NC2=NN=CS2)=O)CCCCCCCCCCCC |
| -1.46 | -0.42 | 1.34 | Masitinib (AB1010) | PDGFR,c-Kit | SelleckChem | S1064 | C1(=C(C=C(C=C1)NC(=O)C2=CC=C(C=C2)CN3CCN(CC3)C)NC4=NC(=CS4)C5=CC=CN=C5)C |
| -1.49 | -0.43 | 0.73 | TAK-733 | MEK | SelleckChem | S2617 | N1(C=NC2=C(C1=O)C(=C(C(N2C)=O)F)NC3=CC=C(C=C3F)I)C[C@@H](O)CO |
| -1.49 | -0.31 | 0.76 | R547 | CDK | SelleckChem | S2688 | N1=C(N=CC(=C1N)C(C2=C(C(=CC=C2OC)F)F)=O)NC3CCN(CC3)S(=O)(C)=O |
| -1.49 | -0.3 | 1.36 | HER2-Inhibitor-1 | HER2,EGFR | SelleckChem | S2752 | N1C=NC(=C2C=C(C=CC=12)C3OC(=CC=3)CNCCS(=O)(=O)C)NC4C=C(C(=CC=4)OC5=CC6N(C=C5)N=CN=6)C |
| -1.49 | -0.54 | 3.69 | PP1 | Src | SelleckChem | S7060 | N1=CN=C2C(=C1N)C(=NN2C(C)(C)C)C3=CC=C(C=C3)C |
| -1.52 | -0.17 | 0.71 | Phenformin HCl | AMPK | SelleckChem | S2542 | C1(=CC=C(C=C1)CCNC(NC(=N)N)=N).Cl |
| -1.53 | -0.79 | 0.76 | KU-0063794 | mTOR | SelleckChem | S1226 | N1=C(N=C2C(=C1N3CCOCC3)C=CC(=N2)C4=CC(=C(C=C4)OC)CO)N5C[C@H](O[C@H](C5)C)C |
| -1.53 | -0.82 | 1.45 | Afatinib (BIBW2992) | EGFR,HER2 | SelleckChem | S1011 | C1(=C(C=C(C=C1)NC2=NC=NC3=C2C=C(C(=C3)O[C@H]4CCOC4)NC(/C=C/CN(C)C)=O)Cl)F |
| -1.6 | -0.76 | 1.57 | HMN-214 | PLK | SelleckChem | S1485 | C1=CC=CC(=C1N(S(C2=CC=C(C=C2)OC)(=O)=O)C(C)=O)/C=C/C3=CC=[N+](C=C3)[O-] |
| -1.65 | -0.65 | 0.85 | Trametinib (GSK1120212) | MEK | SelleckChem | S2673 | C1(N(C(=C2C(=C1C)N(C(N(C2=O)C3CC3)=O)C4=CC(=CC=C4)NC(C)=O)NC5=CC=C(C=C5F)I)C)=O |
| -1.68 | -1.09 | 0.4 | NU7441 (KU-57788) | DNA-PK,PI3K | SelleckChem | S2638 | C1=C(OC2=C(C1=O)C=CC=C2C3=CC=CC4=C3SC5=C4C=CC=C5)N6CCOCC6 |
| -1.73 | -0.39 | 0.78 | TGX-221 | PI3K | SelleckChem | S1169 | C1=C(C=C(C2N1C(C=C(N=2)N3CCOCC3)=O)C(C)NC4=CC=CC=C4)C |
| -1.73 | -0.21 | 0.69 | Asiatic Acid | p38 MAPK | SelleckChem | S2266 | [C@@H]1([C@@H]([C@]([C@]2([C@](C1)([C@]3([C@@](CC2)([C@]4(C(=CC3)[C@@]5([C@@](CC4)(CC[C@H]([C@H]5C)C)C(O)=O)[H])C)C)[H])C)[H])(CO)C)O)O |
| -1.76 | -1.22 | 0.41 | Bardoxolone Methyl | IκB/IKK | SelleckChem | S8078 | C1C(CC[C@@]2(CC[C@]3([C@@]4(CC[C@]5(C(C(C(=C[C@]5(C4=CC([C@]3([C@]2(1)[H])[H])=O)C)C#N)=O)(C)C)[H])C)C)C(=O)OC)(C)C |
| -1.87 | -0.58 | 2.46 | BKM120 (NVP-BKM120, Buparlisib) | PI3K | SelleckChem | S2247 | C1=C(N=CC(=C1C(F)(F)F)C2=NC(=NC(=C2)N3CCOCC3)N4CCOCC4)N |
| -1.87 | -0.81 | -0.28 | WP1066 | JAK | SelleckChem | S2796 | C(/C(=C/C1C=CC=C(N=1)Br)C#N)(=O)N[C@@H](C)C2C=CC=CC=2 |
| -2.47 | -1.81 | 0.47 | Fingolimod (FTY720) HCl | S1P Receptor | SelleckChem | S5002 | C1(=CC=C(C=C1)CCCCCCCC)CCC(CO)(N)CO.Cl |
| -2.81 | -1.04 | -0.68 | Degrasyn (WP1130) | DUB,Bcr-Abl | SelleckChem | S2243 | C1(=CC=CC(=N1)/C=C(/C(N[C@H](C2=CC=CC=C2)CCC)=O)C#N)Br |
| -3.6 | -0.88 | -1.97 | Tideglusib | GSK-3 | SelleckChem | S2823 | S1N(C(N(C1=O)CC2C=CC=CC=2)=O)C3C=CC=C4C=CC=CC=34 |

**Supplementary table 2.** Data collection and refinement statistics for the ULK1 and ULK2 structures.

| **Complex** | **ULK1:PF-03814735** | **ULK2:hesperadin** | **ULK2:MRT67307** | **ULK2:MRT68921** |
| --- | --- | --- | --- | --- |
| **PDB accession code** | **6QAS** | **6QAT** | **6QAU** | **6QAV** |
| **Beamline** | **SLS X06SA** | **SLS X06SA** | **SLS X06DA** | **SLS X06DA** |
| ***Data Collection*** |  |  |  |  |
| Resolution^a^ (Å) | 47.47-1.75 (1.81-1.75) | 48.75-2.77 (2.92-2.77) | 58.50-2.48 (2.61-2.48) | 57.97-2.05 (2.12-2.05) |
| Spacegroup | *P*4_3_2_1_2 | *P*2_1_ | *P*3_1_21 | *P*2_1_ |
| Cell dimensions | *a* = b = 74.2, *c* = 222.3 Å | *a* = 75.2, b = 77.6, *c* = 94.9 Å | *a* = b = 117.0, *c* = 141.5 Å | *a* = 74.7, b = 69.1, *c* = 107.7 Å |
|  | *α, β, γ* = 90.0° | *α, γ* = 90.0˚, *β* = 97.8˚ | *α, β* = 90.0˚, *γ* = 120.0˚ | *α, γ* = 90.0˚, *β* = 97.9˚ |
| No. unique reflections^a^ | 63,838 (6,154) | 27,487 (4,001) | 40,240 (5,786) | 68,365 (6,706) |
| Completeness^a^ (%) | 100.0 (100.0) | 99.2 (99.3) | 100.0 (100.0) | 100.0 (99.9) |
| I/σI^a^ | 14.4 (2.7) | 10.1 (2.0) | 11.5 (2.2) | 11.7 (2.4) |
| R_merge_^a^ (%) | 0.079 (0.806) | 0.071 (0.758) | 0.136 (0.998) | 0.078 (0.657) |
| CC (1/2) | 0.997 (0.777) | 0.997 (0.884) | 0.998 (0.711) | 0.998 (0.842) |
| Redundancy^a^ | 9.2 (9.4) | 5.1 (5.1) | 9.8 (9.8) | 5.3 (5.3) |
| ***Refinement*** |  |  |  |  |
| No. atoms in refinement (P/L/O)^b^ | 4,447/68/589 | 8,141/148/18 | 6,342/102/217 | 8,654/128/430 |
| B factor (P/L/O)^b^ (Å^2^) | 30/33/43 | 109/85/69 | 58/46/52 | 46/27/44 |
| R_fact_ (%) | 17.0 | 21.8 | 19.6 | 19.5 |
| R_free_ (%) | 19.7 | 26.2 | 23.7 | 23.1 |
| rms deviation bond^c^ (Å) | 0.015 | 0.009 | 0.010 | 0.013 |
| rms deviation angle^c^ (°) | 1.5 | 1.1 | 1.1 | 1.3 |
| ***Molprobity Ramachandran*** |  |  |  |  |
| Favour (%) | 96.99 | 95.54 | 96.91 | 97.26 |
| Allowed (%) | 0 | 0 | 0 | 0 |
| Crystallization condition | 2.8 M Ammonium sulfate, 0.1 M citrate, pH 5.6 | 27.5% PEG 3350, 0.1 M sodium citrate, pH 5.9, 0.15 M MgCl_2_, 0.1 M bis-tris, pH 5.75, 5% glycerol | 22.5% PEG 3350, 0.2 M sodium citrate, pH 5.9, 0.1 M bis-tris, pH 5.75, 5% glycerol | 37.5% PEG 3350, 0.1 M sodium citrate, pH 5.9, 0.15 M MgCl_2_, 0.1 M bis-tris, pH 5.5, 5% glycerol |

^a^ Values in brackets show the statistics for the highest resolution shells.

^b^ P/L/O indicate protein, ligand molecules presented in the active sites, and other (water and solvent molecules), respectively.

^c^ rms indicates root-mean-square.
